# An apicomplexan bromodomain, TgBDP1 associates with diverse epigenetic factors to regulate essential transcriptional processes in *Toxoplasma gondii*

**DOI:** 10.1101/2022.12.24.521857

**Authors:** Krista Fleck, Seth McNutt, Feixia Chu, Victoria Jeffers

## Abstract

The protozoan pathogen *Toxoplasma gondii* relies on tight regulation of gene expression to invade and establish infection in its host. The divergent gene regulatory mechanisms of *Toxoplasma* and related apicomplexan pathogens rely heavily on regulators of chromatin structure and histone modifications. The important contribution of histone acetylation for *Toxoplasma* in both acute and chronic infection has been demonstrated, where histone acetylation increases at active gene loci. However, the direct consequences of specific histone acetylation marks and the chromatin pathway that influences transcriptional regulation in response to the modification is unclear. As a reader of lysine acetylation, the bromodomain serves as a mediator between the acetylated histone and transcriptional regulators. Here we show that the bromodomain protein TgBDP1 which is conserved amongst Apicomplexa and within the Alveolata superphylum, is essential for *Toxoplasma* asexual proliferation. Using CUT&TAG we demonstrate that TgBDP1 is recruited to transcriptional start sites of a large proportion of parasite genes. Transcriptional profiling during TgBDP1 knockdown revealed that loss of TgBDP1 leads to major dysregulation of gene expression, implying multiple roles for TgBDP1 in both gene activation and repression. This is supported by interactome analysis of TgBDP1 demonstrating that TgBDP1 forms a core complex with two other bromodomain proteins and an ApiAP2 factor. This core complex appears to interact with other epigenetic factors such as nucleosome remodelling complexes. We conclude that TgBDP1 interacts with diverse epigenetic regulators to exert opposing influences on gene expression in the *Toxoplasma* tachyzoite.

**Summary:** Histone acetylation is critical for proper regulation of gene expression in the single celled eukaryotic pathogen *Toxoplasma gondii*. Bromodomain proteins are “readers” of histone acetylation and may link the modified chromatin to transcription factors. Here, we show that the bromodomain protein TgBDP1 is essential for parasite survival and that loss of TgBDP1 results in global dysregulation of gene expression. TgBDP1 is recruited to the promoter region of a large proportion of parasite genes, forms a core complex with two other bromodomain proteins and interacts with different transcriptional regulatory complexes. We conclude that TgBDP1 is a key factor for sensing specific histone modifications to influence multiple facets of transcriptional regulation in *Toxoplasma gondii*.

## Introduction

The protozoan *Toxoplasma gondii* is a ubiquitous parasite, infecting a third of the world’s human population and vast numbers of livestock and wildlife in sensitive ecosystems. *Toxoplasma* is a member of the phylum *Apicomplexa*, that contains many important pathogens such as the intestinal parasite *Cryptosporidium*, and *Plasmodium* the causative agent of malaria. Infections by this group of intracellular parasites are notoriously difficult to treat or prevent due to their conserved eukaryotic cellular functions and their complex life cycles.

In addition to cell growth and maintenance, rapid molecular changes are required for *Toxoplasma* to transition between life cycle stages to support establishment and persistence of infection. Regulation of chromatin structure to support appropriate gene expression is vital, however these critical mechanisms in *Toxoplasma* are not fully understood (1,2). The additional and removal of post-translational modifications on histones and the importance of this dynamic in regulating transcription has become evident (1,3). Acetylation of lysine residues on histones plays a particularly important role in transcriptional activation (4–10). Similarly to observations in other eukaryotes, abolishing the function of *Toxoplasma* lysine acetyltransferases, or lysine deacetylases, the enzymes responsible for the addition or removal of acetyl groups on histones, perturbs gene expression and disrupts parasite proliferation and life cycle progression (9,11–13). These enzymes have been a focus of investigation for drug discovery, however a key player in the acetylation network, the bromodomain is relatively understudied. The bromodomain consists of an approximately 110 amino acid sequence that forms an alpha helical bundle with a hydrophobic pocket that “reads” (recognizes and binds) the acetyl group on a lysine residue (14). Proteins containing bromodomains may perform multiple functions once bound to their intended targets, largely recruiting and interacting with other complexes to modify chromatin and regulate transcription. Due to their diverse and critical function, and their amenability to small-molecule inhibitors, bromodomains have become promising therapeutic targets in humans as treatments for cancer, immune and metabolic disorders (15).

A handful of bromodomain-containing proteins have recently been identified as key regulators of transcription and potential drug targets in Apicomplexans. In *Plasmodium falciparum* a complex consisting of bromodomain proteins PfBDP1, PfBDP2 and PfBDP7 contributes to the expression of invasion factors (16,17) and in parallel, functions as a repressor complex to maintain mutually exclusive expression of variant surface antigens (VSAs) (18). The bromodomain of a GCN5 homologue in *Toxoplasma* (TgGCN5b) was found to be important for parasite growth and a target of the bromodomain inhibitor L-Moses (11). Another bromodomain inhibitor, IBET-151 was also reported to inhibit *Toxoplasma* tachyzoite proliferation (19). Twelve predicted proteins with conserved bromodomains have been identified in the *Toxoplasma* genome, but six of these (TgBDP1-6) are unique to early branching eukaryotes (20,21). While these parasite-specific bromodomain proteins have excellent therapeutic potential, they have yet to be studied in *Toxoplasma* and their functions are unknown.

In the present study we sought to determine the role of the parasite-specific bromodomain protein TgBDP1 in *Toxoplasma* tachyzoites and validate its potential as a therapeutic target. Through sequence and structure analysis we confirmed TgBDP1 has a conserved bromodomain with homologues displaying a similar domain architecture only present in early branching alveolates. We generated a tetracycline-regulatable *tgbdp1* knockdown line and show that TgBDP1 is essential for progression through the parasite lytic cycle. We adapted and performed the Cleavage Under Targets & Tagmentation (CUT&Tag) technique for the first time in a protozoan, revealing that TgBDP1 is recruited to transcriptional start sites and performed co-immunoprecipitations to identify interacting proteins. Further supporting its role in transcription, knockdown resulted in substantial disruption to parasite gene expression. We conclude that TgBDP1 is an essential component of several key transcriptional regulatory complexes.

## Results

### TgBDP1 is a bromodomain-containing protein conserved in apicomplexans

The *Toxoplasma* genome encodes twelve bromodomain-containing proteins, six of which (named TgBDP1-TgBDP6), have no homologues in mammals, plants, or fungi (20,21). TgBDP1 (TGME49_263580) is predicted to be a 76 kDa protein containing a single bromodomain and a series of three ankyrin repeats (Fig. 1A). This domain architecture is only found in proteins from a subset of alveolates. BLAST analyses and conserved domain searches identified homologues in all Apicomplexan species examined but only a handful of other alveolates (Fig. 1B-C). The bromodomain is highly conserved and contains the characteristic tyrosine (Y) and asparagine (N) residues necessary for domain binding to acetylated lysines (Fig. 1D) (22). The structure of the bromodomain of TgBDP1 was modelled using I-TASSER and compared to experimentally determined bromodomain structures (23). The predicted structure of TgBDP1 is highly similar to that of the bromodomain from the human protein BAZ2B, forming an alpha helical bundle and hydrophobic binding pocket in which the residues required for coordination of the target acetylated lysine are present and appropriately positioned (Fig. 1E). The crystal structure of the bromodomain of *Plasmodium falciparum* PfBDP1 has recently been determined (PDB: 7M97) (24) and closely matches that of the predicted structure of the bromodomain in TgBDP1. The conserved sequence and structure of the predicted TgBDP1 bromodomain suggests that TgBDP1 is a functional acetyllysine reader.

**Figure 1.**
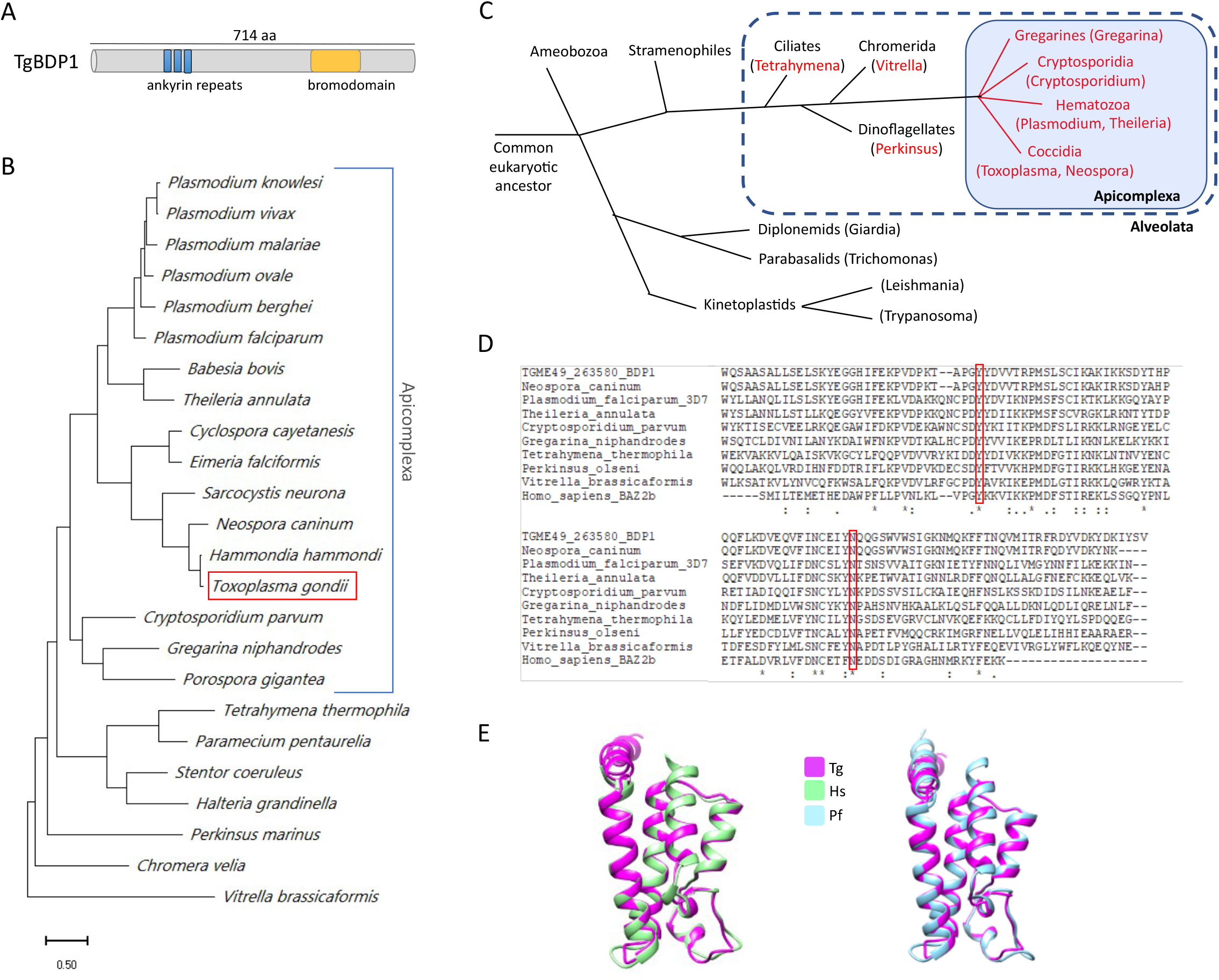
TgBDP1 is a bromodomain containing protein that is conserved among alveolates. A) Depiction of TgBDP1 protein size and domain architecture. B) Phylogenetic tree of TgBDP1 protein homologues drawn to scale, with branch lengths measured in the number of substitutions per site. C) Evolutionary tree with genera containing predicted TgBDP1 homologues in red. Apicomplexans are in blue shaded box and blue dotted line encompasses alveolates. Branch lengths are not to scale. D) Multiple alignment of bromodomain amino acid sequences from representative species, with TgBDP1 denoted as TGME49_263580 BDP1. The highly conserved tyrosine (Y) and asparagine (N) residues required for binding acetylated lysines are boxed in red. E) The predicted structure of the TgBDP1 bromodomain (pink) overlaid with the *Homo sapiens* B2AZB bromodomain (green, PDB:5DYU) and *Plasmodium falciparum* PfBDP1 bromodomain (blue, PDB:7M97).

### TgBDP1 has an mRNA isoform

The predicted protein sequence of TgBDP1 contains 714 amino acids. However, intron predictions based on available RNA-sequencing data in the *Toxoplasma* genome database ToxoDB (25) predicted two possible sizes for the first exon (Supplemental Fig. 1A). The transcript with the longer exon matches the predicted mRNA sequence for *tgbdp1*, and the transcript containing the shorter exon produces an isoform with 63 fewer nucleotides, equivalent to a loss of 21 amino acids (Supplemental Fig. 1B). High resolution nanopore sequencing of *Toxoplasma* mRNAs conducted by Lee et al. also detected the two isoforms (Supplemental Fig. 1A) (26). To confirm these findings, we amplified and sequenced *tgbdp1* cDNA. Half of the clones sequenced (3/6) contained the shorter isoform, which we named *tgbdp1a* (Supplemental Fig. 1C). Additionally, our RNA-sequencing data from a separate line of experiments (described later) showed *tgbdp1* peak variations in all three replicates that would be consistent with a mixed population of the two mRNA isoforms (Supplemental Fig. 1D). The TgBDP1 and TgBDP1a proteins are predicted to be 76 kDa and 74.5 kDa, respectively. However, this small difference in size cannot be distinguished by Western blotting of N-terminally or C-terminally tagged TgBDP1 (Fig. 2D, Fig. 4A). The remaining experiments outlined in this study are focused on TgBDP1 that was modified at the endogenous genomic locus, and thus TgBDP1 isoforms would presumably be processed and function as normal. Additional studies will be needed to determine if these two isoforms have distinct functions.

**Figure 2.**
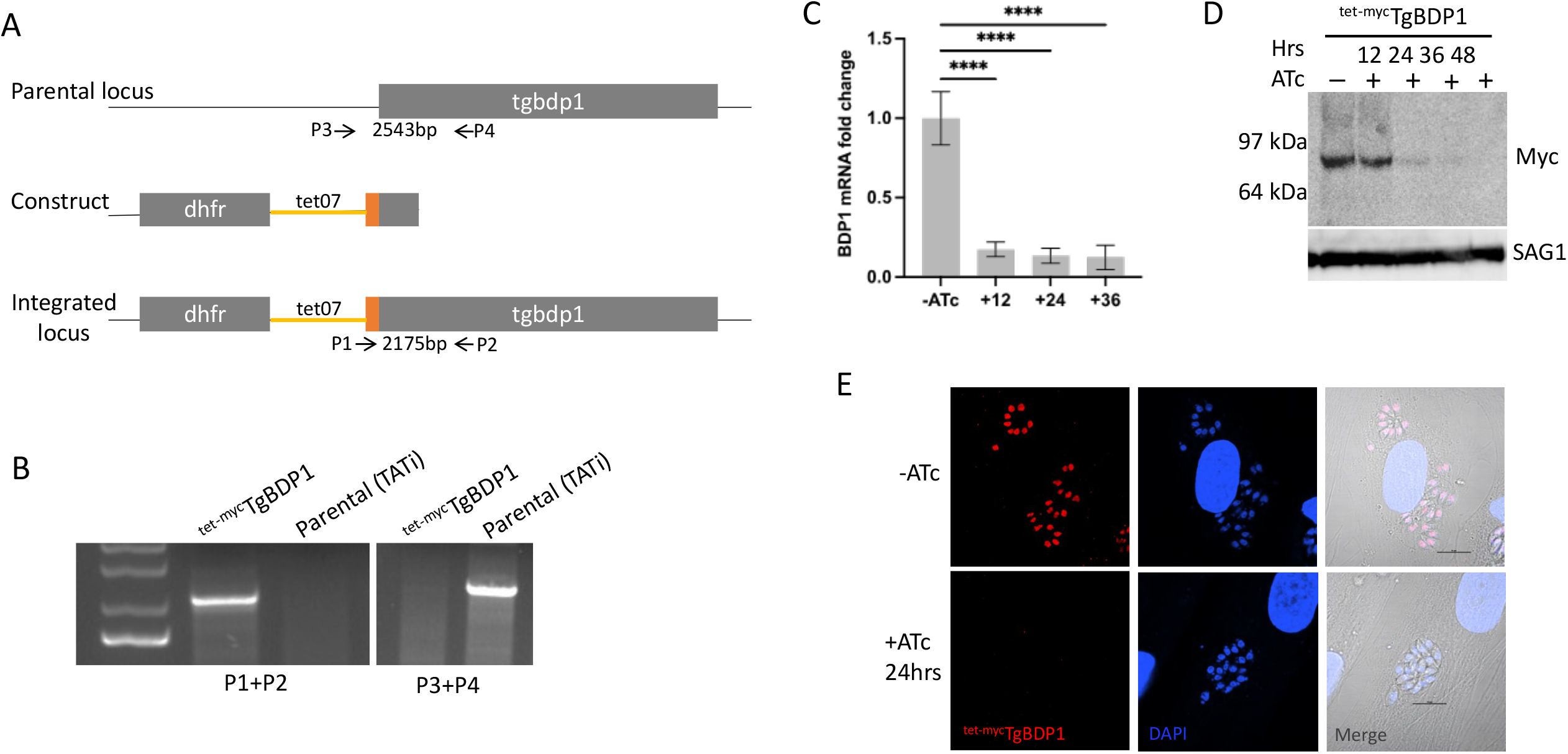
Generation of a *tgbdp1* inducible knockdown. A) Strategy for *tgbdp1* promoter replacement with a tetracycline-regulatable promoter, and insertion of an N-terminal myc tag (orange). The *dhfr* gene was inserted for selection of transgenic parasites with pyrimethamine resistance. Primers used to confirm integration are included. B) PCRs confirming correct genomic modification. Primers P1 and P2 amplify a 2,175bp fragment only present in the transgenic line (^tet-myc^TgBDP1), and primers P3 and P4 amplify a 2,543bp fragment only in the parental (TATi) genome. C) RT-qPCR of *tgbdp1* mRNA levels normalized to the -ATc sample, n=3, **** = p-value <0.0001. D) Western blotting of ^tet-myc^TgBDP1 lysates from parasites cultured -ATc and +ATc for 12, 24, 36, and 48hrs. TgSAG1 was included as a loading control. E) IFA of ^tet-myc^TgBDP1 parasites cultured 24hrs -ATc and +ATc.

### Generation of TgBDP1 inducible knockdown

A genome-wide CRISPR screen to evaluate essentiality of genes in *Toxoplasma* tachyzoites reported a low fitness score for *tgbdp1*, indicating that this gene could be essential (27). To determine the role of TgBDP1, we generated a transgenic, inducible knockdown line ^tet-myc^TgBDP1 in which the endogenous *tgbdp1* promoter is replaced with a hybrid of the *Toxoplasma sag4* promoter and the tetracycline-responsive promoter, and a sequence encoding a triple myc tag is inserted at the 5’end of the gene (Fig. 2A). Correct genomic integration was confirmed by PCR (Fig. 2B), and nuclear localization of TgBDP1 protein was determined by immunofluorescence (IFA) (Fig. 2E). Treatment with anhydrotetracycline (ATc) to abolish transcription of *tgbdp1* results in a significant reduction of *tgbdp1* mRNA levels (Fig. 2C). TgBDP1 protein also decreased over time as seen by Western blotting and IFA, falling below detectable levels by 36 hours (Fig. 2D-E).

### TgBDP1 is essential for the tachyzoite lytic cycle

Plaque assays were performed to evaluate parasite proliferation in the absence of TgBDP1. Tachyzoites of ^tet-myc^TgBDP1 did not form plaques in the presence of ATc over six days, unlike the parental parasite line (TATi) that grew normally and formed plaques in the presence and absence of ATc (Fig. 4A). To determine the precise point in the parasite’s lytic cycle that is inhibited, we tested the ability of parasites to invade and replicate within host cells. Parasites exposed to ATc for 36hrs displayed a three-fold reduction in their ability to invade cells (Fig. 4B). Defects in replication were seen as early as 12hrs post-invasion in the presence of ATc (Fig. 4C). By 36hrs parasites were significantly deformed and completely stalled in their replication. No defects in replication were observed when the parental line (TATi) was exposed to ATc (Supplemental Figure 2). These results demonstrate that loss of TgBDP1 severely impedes tachyzoite host cell invasion and replication, and TgBDP1 is essential for parasite proliferation.

### TgBDP1 functions as part of a parasite-specific core complex

Bromodomain-containing proteins generally function as a component of a larger epigenetic regulatory complex. We sought to determine the function of TgBDP1 during tachyzoite proliferation by identifying TgBDP1-interacting proteins. A transgenic parasite line was generated in which the C-terminus of TgBDP1 was tagged with a 3xHA epitope tag. Western blotting and IFA confirmed the correct protein size (81kDa) and nuclear localization of TgBDP1 (Fig 4A). This parasite line was used for co-immunoprecipitations (coIPs) followed by mass spectrometry of the pulldown to identify TgBDP1-associated proteins.

Peptide counts from three independent replicates were submitted to REPRINT (https://reprint-apms.org/) (28) for SAINT analysis to identify the most highly significant proteins that interact with TgBDP1 (Fig 4B). SAINT analysis evaluates total peptide counts between control and test samples and assigns a score (between 0 and 1) to each protein hit that describes the probability that a protein is a genuine interactor of TgBDP1. The most significant and abundant protein isolated alongside TgBDP1 with a SAINT score of 1, was the bromodomain protein TgBDP2 (Fig. 5B). Two other proteins were also assigned SAINT scores of 1, including the bromodomain-containing protein, TgBDP5, and the AP2 factor TgAP2VIIa-7 that contains a PHD and a SET domain. In addition to this core network of factors, 10 additional proteins with predicted nuclear localization were identified as potentially significant interactors (Fig. 4B, Supplemental Table 1), including predicted transcription factors, chromatin remodeling factors, RNA-associated proteins and enzymes involved in DNA, RNA and chromatin-related pathways (Fig 4B).

The significant enrichment, but low abundance and diverse function of these TgBDP1-bound proteins suggests that TgBDP1 functions as a component of multiple complexes involved in a variety of chromatin-related processes. These findings imply that TgBDP1 interacts with TgBDP2, together with TgBDP5 and TgAP2VIIa-7 to function as part of multiple functional complexes.

### TgBDP1 is found at transcriptional start sites of active genes

We next addressed the occupancy of TgBDP1 across the *Toxoplasma* genome by using Cleavage Under Targets & Tagmentation (CUT&Tag). This technique has several advantages over traditional ChIP-seq and has provided high quality results with human, mouse, and zebrafish cells (29). We adapted the technique for use with *Toxoplasma* parasites and used our transgenic TgBDP1^HA^ line to determine TgBDP1 genome-wide localization. Briefly, an anti-HA antibody was used to target transposases harboring sequencing tags to chromatin bound TgBDP1. The transposase then cleaved DNA on either side of the TgBDP1 binding site, adding sequencing tags to the DNA ends. Tagged DNA fragments were amplified, sequenced, and mapped to the *Toxoplasma* genome. Our results were consistent between three independent replicates with little to no signal in the parental negative controls (Fig. 5A). To validate the CUT&Tag approach in *Toxoplasma*, we also performed the experiment with an antibody to acetylated lysine 9 on histone H3 (H3K9ac), a well-known active gene marker that is often found at transcription start sites (TSS). The distribution of H3K9ac observed by CUT&Tag was consistent with previous ChIP-chip studies (30). An average of approximately 5,400 peaks for TgBDP1 binding sites were identified, the majority of which (63%) were located upstream of protein-coding genes (Fig. 5B-C). Only 39% of TgBDP1 peaks located at transcriptional start sites coincided with H3K9ac peaks (Fig. 5D). These results suggest that TgBDP1 may bind or coincide with H3K9ac, but not exclusively; it likely has roles independent of this particular acetyl mark. Our observation that TgBDP1 was bound upstream of open reading frames prompted us to compare TgBDP1 binding sites to predicted transcriptional start sites obtained from a recent study by Markus et al. (31). Of the 2,155 TgBDP1 peaks found within 2kb upstream of protein-coding genes, a large proportion (42%) align directly with transcriptional start sites (Fig. 5C-D, Supplemental Table 2).

The expression profiles of TgBDP1 target genes were compared between tachyzoites and parasites undergoing bradyzoite differentiation. Using previously published transcriptomic data from Waldman et al. (32), we plotted the relative abundance of transcripts of TgBDP1-target genes during tachyzoite replication (X-axis) and early bradyzoite differentiation (Y-axis) (Fig 6A). Most TgBDP1-target genes are consistently expressed under both growth conditions (grey markers) suggesting that TgBDP1 is recruited to constitutively active genes. We also observed TgBDP1 binding to promoters of genes that are induced during bradyzoite formation (blue markers), suggesting that TgBDP1 may play a role in regulating chromatin structure to “poise” a gene for increased expression in response to environmental signals. However, a subset of TgBDP1-target genes are expressed at a low, or undetectable level in both tachzyoites and early bradyzoites (lower left side of the plot), demonstrating that TgBDP1 binding within a gene promoter does not directly correlate with active or poised transcription and indeed, TgBDP1 may act as a repressor on these genes.

Gene Ontology (GO) enrichment analysis was performed on TgBDP1 target genes, to determine if it is a regulator of specific functional pathways. TgBDP1 was bound to the promoter of 24% of all annotated *Toxoplasma* genes so accordingly, functional enrichment analysis identified a diverse range of different biological process that may be subject to TgBDP1 regulation. Those pathways that were significantly enriched include transcription and mRNA splicing, ribosomal formation and protein maturation, and metabolic processes associated with the mitochondrion. Manual inspection of the list of TgBDP1 target genes also revealed many transcriptional regulators that may be subject to TgBDP1 regulation. Of the 85 predicted transcription factors in the *Toxoplasma* genome, 33% were found to have TgBDP1 at their TSS, including 25 ApiAP2s and the myb regulator of bradyzoite formation, *tgbfd1* (Fig. 6B) (32). Supporting our observation that *tgbdp1*-knockdown parasites were defective in host cell invasion, 105 genes encoding microneme, rhoptry and dense granule proteins are putative TgBDP1 targets.

TgBDP1 is recruited to many, but not all active genes in the tachyzoite, so we performed motif enrichment analysis of TgBDP1-target gene promoters to identify any specific DNA sequence motifs that may contribute to TgBDP1 recruitment. MEME-ChIP analysis of sequence 250bp up- and down-stream of the predicted TSS identified two DNA sequence features that were significantly enriched in target promoters (Fig 6C). The motif GCATGCA (motif 1) and a degenerate pyrimidine-rich sequence (motif 2) that is enriched downstream of the TSS of selected genes. Both of these motifs have previously been reported as characteristics of *Toxoplasma* TSSs (31,33–35).

### Loss of TgBDP1 impacts parasite transcription

Loss of TgBDP1 caused a defect in host cell invasion and an arrest in tachyzoite replication. As TgBDP1 appears to be a component of at least one epigenetic complex and is recruited to approximately 25% of predicted open reading frames, we investigated if the observed growth arrest during knockdown was due to global dysregulation of parasite transcription. Western blotting analysis of TgBDP1 during knockdown (Fig. 2D) indicates that TgBDP1 protein levels fall below 50% by 24hr and are almost undetectable by 36hr. To determine the essential contribution of TgBDP1 to transcriptional regulation, we performed RNA-seq at 12, 24 and 36hr after addition of ATc. We observed a time dependent effect of *tgbdp1* knockdown on the parasite transcriptome with a small number of genes impacted at 12 hours, but larger impacts observed at later time points (Fig. 7A). More than half of differentially expressed genes identified at 24hr were also dysregulated at 36hr (Fig. 7B). This observation, in addition to phenotypic data showing complete growth arrest at 36hr (Fig. 3), suggests that differentially expressed genes at 12 and 24hr likely represent more direct effects of *tgbdp1* knockdown, but by 36hr there are more indirect effects on parasite gene expression. Therefore, we focused our analyses on differentially expressed genes at the 12 and 24hr timepoints.

**Figure 3.**
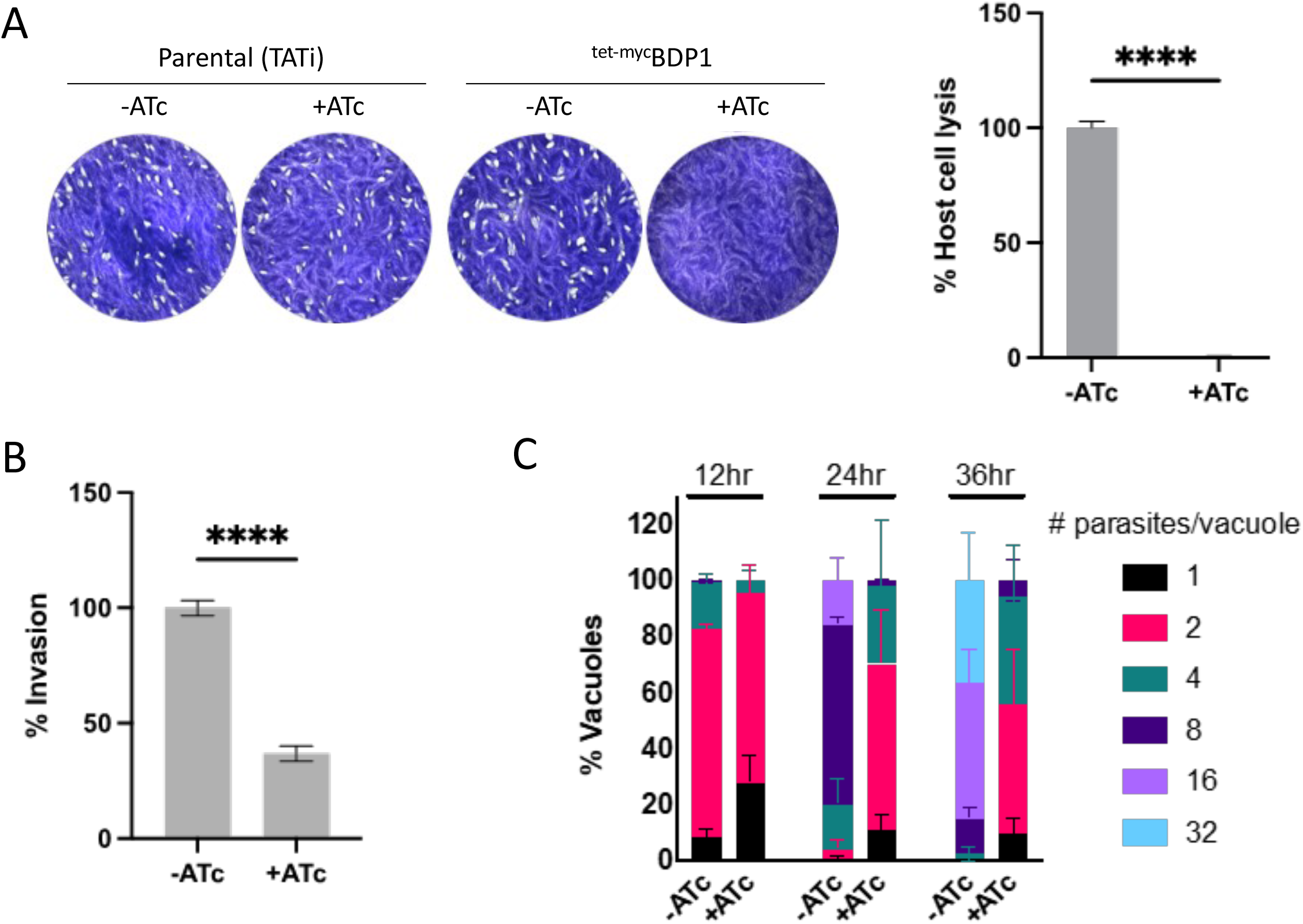
Loss of TgBDP1 causes significant defects in host cell invasion and parasite replication. A) Plaque assays were conducted with the ^tet-myc^TgBDP1 and parental (TATi) parasite lines -ATc and +ATc. Images are representative from three independent experiments after six days of growth. The area lysed was calculated as a percentage of -ATc. B) Invasion assays were performed by counting the number of parasites that invaded host cells and calculated as a percentage of -ATc. C) Doubling assays were performed by counting the number of parasites per vacuole for 100 vacuoles at 12, 24 and 36hrs after inoculation, and plotted as a percentage of the total number of vacuoles. All experiments were done in triplicate and unpaired t-tests performed for plaque and invasion assays, n=3, **** = p-value <0.0001.

**Figure 4.**
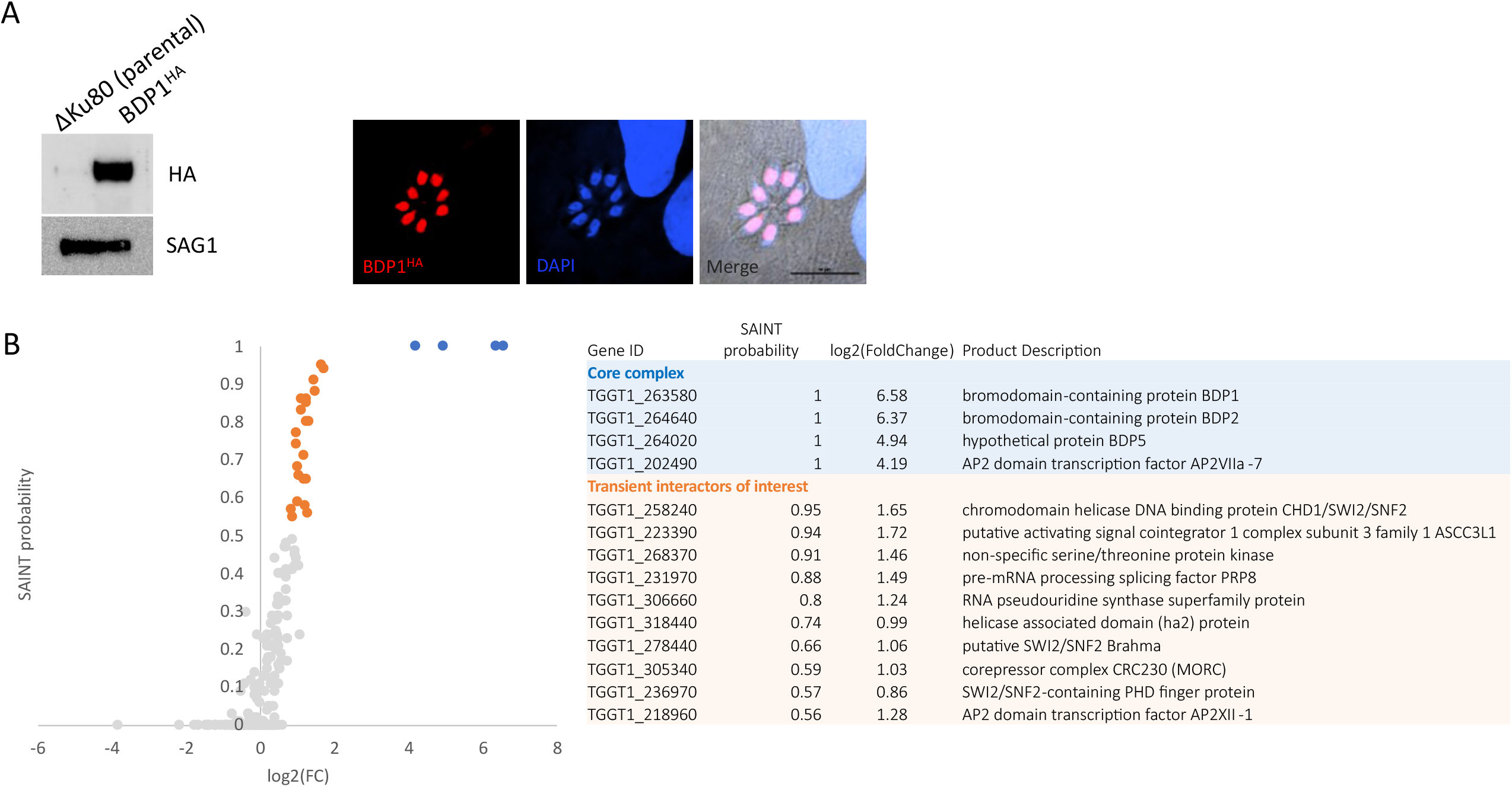
TgBDP1 is a nuclear protein that interacts with transcriptional and chromatin regulatory proteins. A) TgBDP1 was tagged at the C-terminus with 3xHA. Western blotting of parasite lysate showed the tagged protein at the predicted size (81kDa) with no signal detected in the parental line (Δku80). SAG1 was used as a loading control. Nuclear localization of tagged TgBDP1 was confirmed by IFA. B) CoIPs of both TgBDP1^HA^ and parental lines were conducted and enriched proteins identified by mass spectrometry. SAINT probability scores are plotted against log2(fold change) from all three replicates. TgBDP1 and its most significant interactors are plotted in blue. Other highly probable interactors are plotted in orange. Scores and gene IDs are detailed in the adjacent table.

**Figure 5.**
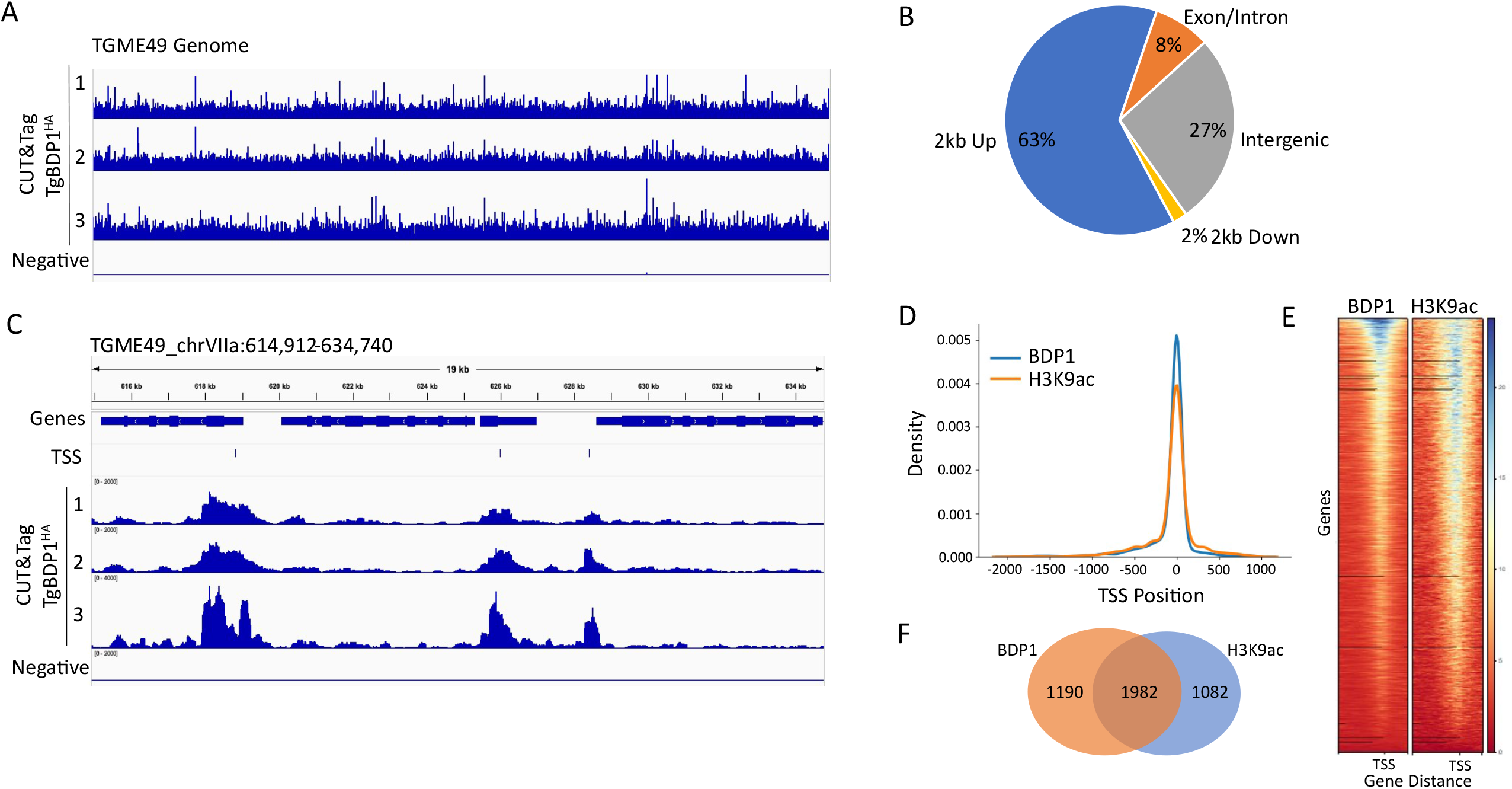
TgBDP1 binds upstream of many protein coding genes. A) Alignment of peak intensities of three replicates of TgBDP1 CUT&Tag and a negative control mapped to the *Toxoplasma* ME49 genome. The data range is set the same between all four tracks. B) A breakdown of the location of all TgBDP1 peaks relative to protein-coding genes. C) Representative snapshot of TgBDP1 peaks aligning with transcriptional start sites (TSS) of three genes on chromosome VIIa. D) Density graph of TgBDP1 and H3K9ac peaks located - 2kb and +1kb from TSS. E) Heatmap of TgBDP1 and H3K9ac peak densities at gene distances from TSS. F) Venn diagram of the number of genes with TgBDP1 and H3K9ac peaks at TSS.

A small number of genes were significantly dysregulated by two-fold or more at 12hr. Most significant, was upregulation of *tgbdp2*, the gene encoding TgBDP2, the binding partner of TgBDP1 (Fig. 7C, Supplemental Table 3). Of the 48 significantly downregulated genes, TgBDP1 is recruited to the TSS of only four – ROP36 (TGME49_207610), an aspartyl protease expressed in sporulated oocysts (TGME49_272510) (36), a Toxoplasma gondii family D protein (TGME49_313000) and a hypothetical protein (TGME49_243700). Apart from *tgbdp1* itself, only one of the genes that are significantly downregulated has a phenotype score < -1 (TGME49_211000), therefore downregulation of these genes is unlikely to contribute directly to parasite growth arrest. At this early timepoint in *tgbdp1* downregulation, there is very little impact on gene expression, aside from the significant upregulation of *tgbdp2* which is likely a compensatory response to loss of TgBDP1.

A more dramatic impact on parasite transcription was observed by 24hr post *tgbdp1* knockdown. A total of 550 genes were downregulated two-fold or more, only 118 of which are bound by TgBDP1 at their TSS (Fig. 7C, Supplemental Table 3). Gene Ontology (GO) Enrichment Analysis was performed on the lists of significantly dysregulated genes but did not identify any specific functional pathways that were enriched. Unexpectedly, a larger number of genes were upregulated in response to *tgbdp1* knockdown. Of the 675 upregulated genes, 140 are also bound by TgBDP1 at the TSS. GO enrichment analysis of upregulated genes identified over four-fold (p=0.01) enrichment of genes encoding surface proteins and invasion-related proteins from the SAG, microneme, dense granule and rhoptry families. Although 105 invasion genes were identified by CUT&Tag to be associated with TgBDP1, only 19 of those were upregulated. Therefore, the majority of dysregulated invasion genes are likely indirectly regulated by TgBDP1. These results suggest that loss of TgBDP1 influences the expression of other critical transcriptional activators or repressors; TgBDP1 is bound to the TSS of many putative transcription factors (Fig. 6B) and several transcription factors are dysregulated during *tgbdp1* knockdown (Fig. 7C).

**Figure 6.**
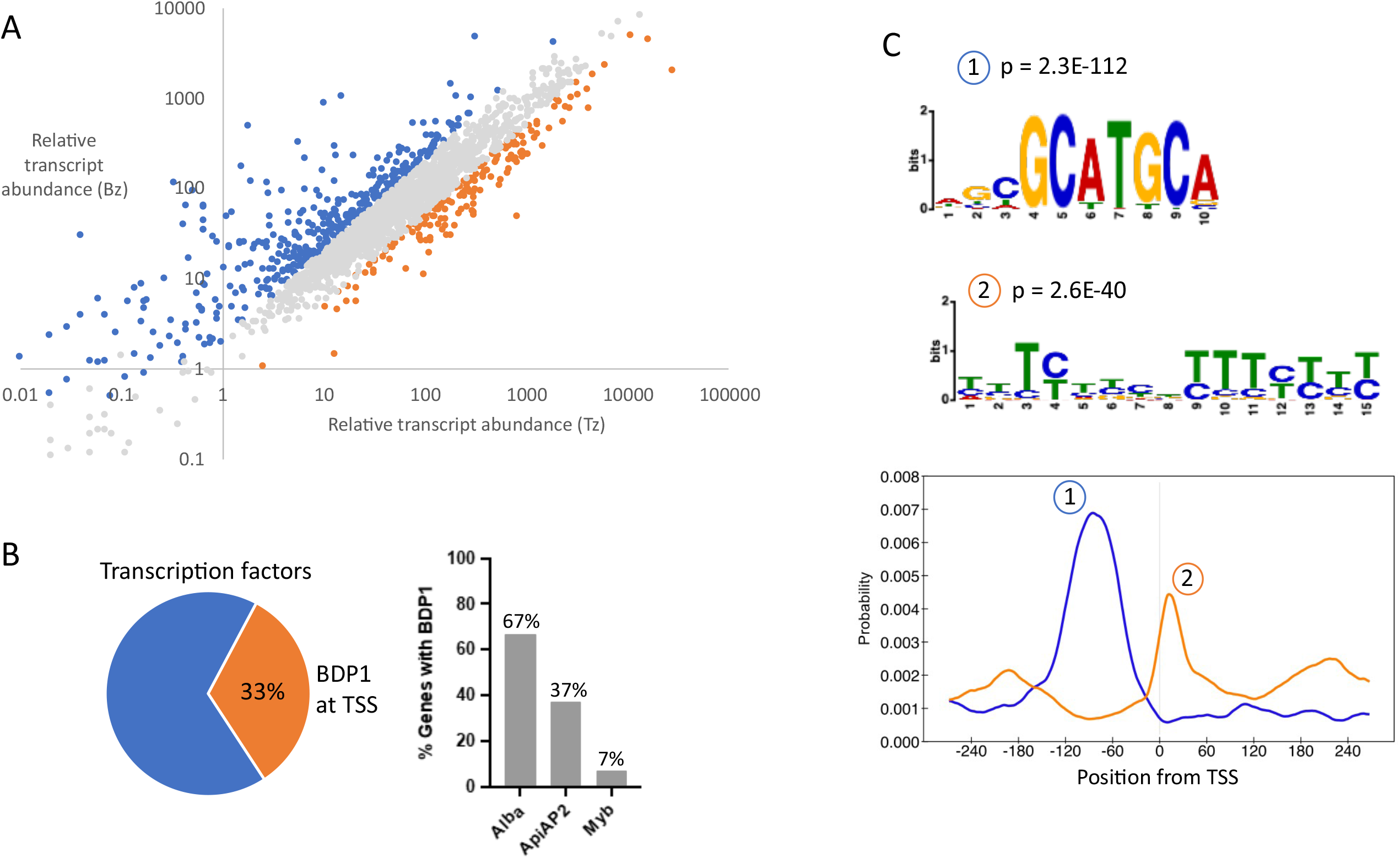
TgBDP1 is predominantly recruited to promoters of active genes. A) Relative transcript abundance of putative TgBDP1 target genes in tachyzoites (x-axis) and after 48hr of bradyzoite induction (y-axis) (32). Blue markers: transcripts upregulated two-fold or more; Orange markers: transcripts downregulated two-fold or more; Grey: transcripts not significantly changed during bradyzoite differentiation. B) Left: pie chart of percentage of predicted transcription factor genes with (orange) and without (blue) TgBDP1 bound. Right: bar graph showing percentage of each family of transcription factor genes bound by TgBDP1. C) Motif analysis of TSS sequences associated with TgBDP1. Top: the two most significant motifs identified and their p-values. Bottom: location of each motif from TSS.

**Figure 7.**
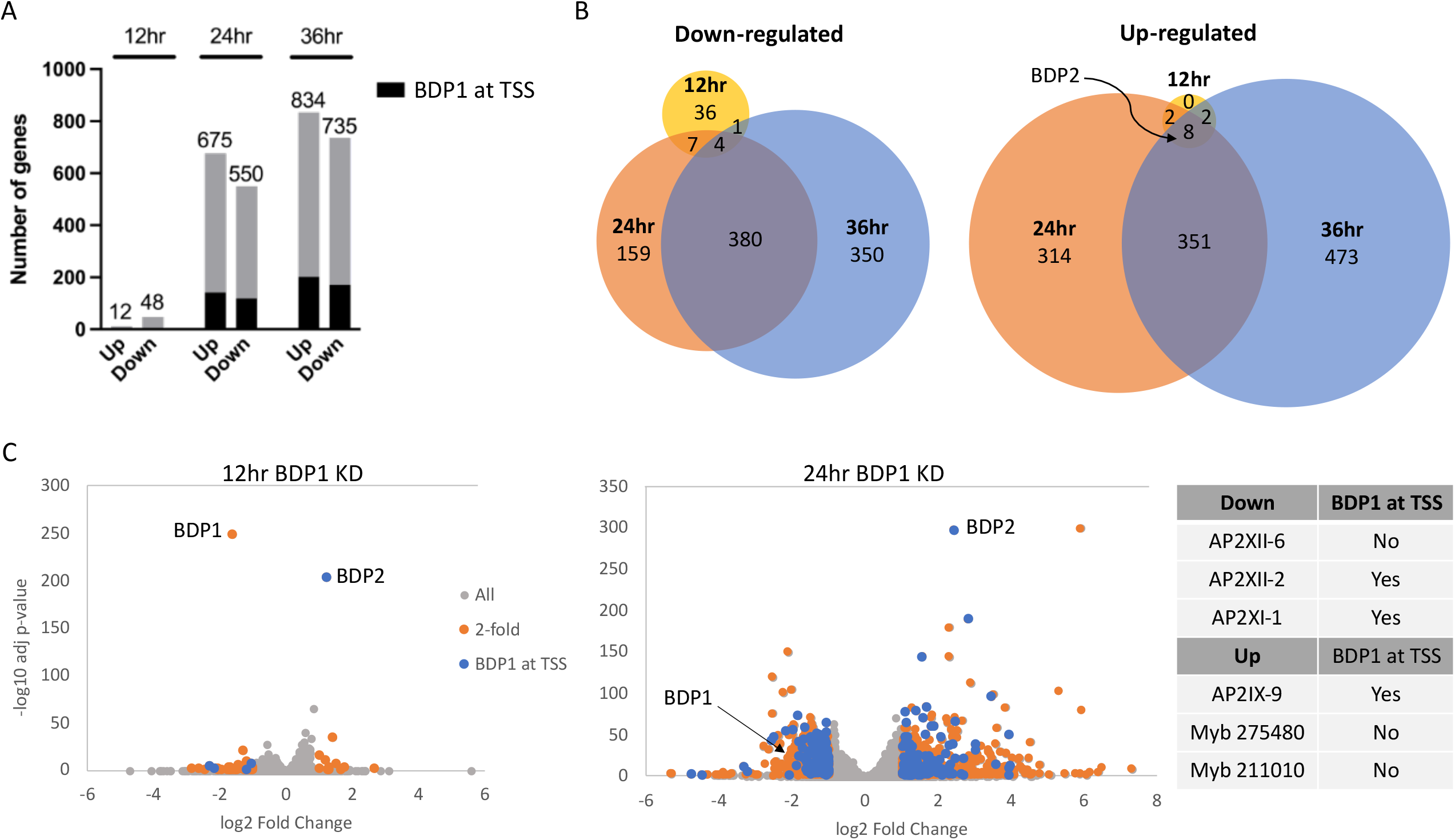
TgBDP1 down-regulation causes global dysregulation of gene expression. A) Bar graph depicting the number of genes up- and down-regulated identified by RNA-seq in ^tet-myc^TgBDP1 parasites incubated for 12, 24 and 36hrs with ATc. Black shading indicates the number of those genes with TgBDP1 found at the gene TSS from CUT&Tag analysis of TgBDP1 binding sites. B) Venn diagrams of genes up- and down-regulated between all three timepoints. TgBDP2 is the only gene differentially expressed at all three timepoints that is also bound by TgBDP1 at its TSS. C) Volcano plots of differentially genes in ^tet-myc^TgBDP1 knockdown parasites at 12 and 24hrs. Genes up- or down-regulated two-fold or more are in orange and those also identified as TgBDP1-bound by CUT&Tag are in blue. The table shows specific transcription factors differentially expressed at 24hr post knockdown.

The most consistent effect of *tgbdp1* knockdown is sustained upregulation of *tgbdp2*. This is the only gene identified at all three timepoints that was differentially expressed that also had TgBDP1 bound at the TSS, per CUT&Tag analysis (Fig. 7B). It remains unclear whether residual TgBDP1 is directly involved in upregulating *tgbdp2* expression or if loss of TgBDP1 triggers an indirect compensatory response. Overall, the large number and functional diversity of both up- and down-regulated genes impacted during *tgbdp1* knockdown supports a global chromatin regulation function for TgBDP1, rather than merely a transcriptional activator. Furthermore, the subset of transcription factors that are directly and indirectly impacted by TgBDP1 points to this bromodomain and its complex(es) as key players of *Toxoplasma* gene expression.

## Discussion

Histone acetylation is associated with activation of gene expression. Specific histone acetylation marks typical for gene activation, such as H3K9Ac and H4K8Ac, K12Ac and K16Ac are enriched at active gene promoters in *Toxoplasma* tachyzoites (8,10,30). In addition to altering chromatin structure to facilitate transcription, histone acetylation serves to recruit regulatory complexes to specific loci through the action of reader modules, such as bromodomains, although the precise function of most bromodomain-containing proteins in *Toxoplasma* gene regulation has yet to be determined. To understand the contribution of bromodomain proteins in *Toxoplasma* in mediating signals between histone acetylation marks and transcription initiation complexes, we investigated the role of a protein unique to alveolates and conserved within the Apicomplexa, TgBDP1. Based on reports on the *Plasmodium falciparum* BDP1 homologue (16,17), we initially hypothesized that TgBDP1 was a transcriptional activator that binds to activating acetylation marks on histone tails to recruit AP2 proteins and other transcription factors to active gene promoters. This was supported by both our CUT&Tag and proteomic analysis that found TgBDP1 bound to chromatin at the predicted TSS of many active genes in tachyzoites and associated with putative transcription factors and epigenetic regulators. However, analysis of global transcription revealed that more genes were upregulated than downregulated during ablation of TgBDP1, indicating a more complex role for this regulatory factor. Furthermore, we did not find a correlation between dysregulated genes and those that have TgBDP1 bound at the TSS, suggesting that much of the impact on gene expression that we observed is due to indirect effects of *tgbdp1* knockdown. TgBDP1 may influence transcription through regulation of transcriptional activators and repressors or via other, non-chromatin binding events, such as associating with acetylated transcription machinery.

Proteomic analysis of the TgBDP1 interactome supports multiple regulatory functions for the core TgBDP1/2/5 complex. We identified consistent association between the three bromodomain-containing proteins and the AP2 factor AP2VIIa-7, which contains a methyltransferase domain in addition to the AP2 domain. Our proteomic approach also detected transient interactions between TgBDP1 and three SWI/SNF nucleosome remodelers, implying a role for this complex in regulation of chromatin structure. Modulation of local chromatin structure can serve to both facilitate or repress transcription. TgBDP1 acting as a major regulator of chromatin structure would explain its apparent contrary influence on gene expression. We should also consider the presence of two isoforms of TgBDP1 (Fig S1) that may define different complex compositions and functionalities. Our analysis also detected an interaction between TgBDP1 and TgMORC, one of the major regulators of parasite stage-specific gene expression. TgMORC is required for recruitment of the HDAC3 repressor complex to maintain silencing of specific genes during the tachyzoite stage (37). The association of TgBDP1 with TgMORC hints at a role for TgBDP1 in gene silencing and may explain the large number of upregulated genes during *tgbdp1* knockdown (almost half of the genes (307 out of 676) upregulated during *tgbdp1* knockdown are TgMORC target genes), but further study of this interaction is necessary to understand the functional relevance to parasite transcriptional regulation. TgMORC transcript levels were also slightly increased during *tgbdp1* knockdown (1.56-fold) but were excluded by our cut offs. If this increase in TgMORC transcript levels results in functionally relevant increase in TgMORC protein, then this may additionally explain the downregulation of genes that do not have TgBDP1 bound at their TSS. Another interactor with the TgBDP1 core complex was TGGT1_235420, a protein of unknown function that localizes to the nucleus. Although the function of this protein is unknown, it is essential for tachyzoite survival in both *Toxoplasma* and the related coccidian *Neospora caninum* (38)

Many of the genes that are upregulated in response to *tgbdp1* knockdown are more highly expressed in bradyzoites or sexual stages compared to tachyzoites (Fig. S3). It is unclear if TgBDP1 is directly repressing expression of these genes, or if it indirectly represses stage-specific genes by regulating transcriptional factors required for maintenance of appropriate stage expression. Transcription factors dysregulated during *tgbdp1* knockdown include three AP2 factors that are significantly downregulated (AP2XI-1, AP2XII-2 and AP2XII-6). One of these, AP2XII-2 is a cell cycle-regulated protein associated with the TgMORC/HDAC3 repressor complex (37). Sustained AP2XII-2 expression may be required during tachyzoite growth to target the MORC complex to sexual or bradyzoite stage genes and maintain tachyzoite proliferation. Indeed, Srivastava and colleagues demonstrated that knockdown of AP2XII-2 results in slowed tachyzoite replication and increased cyst formation (39)(Srivastava 2022). We also observed significant upregulation of genes encoding three putative DNA binding proteins: two myb domain-containing proteins and the AP2 factor AP2IX-9. The functions of the two myb domain proteins are unknown, however they are both consistently expressed across all life cycles stages, suggesting that they may have housekeeping functions, and their upregulation may be a compensatory response to loss of TgBDP1. AP2IX-9 is a repressor of bradyzoite commitment, that is normally upregulated in response to bradyzoite induction conditions (39). TgBDP1 is bound to the TSS of AP2IX-9 in unstressed tachyzoites, so it is unclear how loss of TgBDP1 contributes to upregulation of AP2IX-9. There are a couple of possibilities for TgBDP1’s influence on *ap2ix-9* expression; TgBDP1 maintains repression of *ap2ix-*9 directly through its interactions with TgMORC and that loss of TgBDP1 derepresses *ap2ix-9*. Alternatively, TgBDP1 poises *ap2ix-9* for expression and upregulation of *ap2ix-9* is a direct response to the stress induced in the parasites by loss of TgBDP1. Another major regulator of parasite stage transition TgBFD1, which promotes upregulation of bradyzoite genes during the initial stages of differentiation into tissue cysts was also slightly upregulated (1.83-fold). However, since TgBFD1 appears to be primarily regulated at the translational level, and the protein that drives translation of TgBFD1, TgBFD2 (40) is not significantly increased during *tgbdp1* knockdown, it is unclear if this slight increase in *tgbfd1* transcript leads to increased protein levels.

One family of proteins upregulated during *tgbdp1* knockdown are the SRS family of surface proteins. This mirrors a recent report that evaluated gene expression patterns during knockdown of TgBDP5, a bromodomain protein that is also a component of the TgBDP1 core complex (42). Many of those upregulated during *tgbdp1* knockdown peak in expression level during sexual stages or the bradyzoite tissue cyst (41). A single cell RNA-seq analysis revealed that during normal tachyzoite growth, expression of many “stage-specific” SRS surface antigens is variable between individuals within a population of clonally derived parasites (43). Although the function of these surface proteins is unknown, the authors of these studies speculate that they may contribute to antigenic variation. In *P. falciparum*, many of the variable surface antigens *var, sica* and *rifin* were derepressed during knockdown of either the BDP1 (PfBDP1) or BDP5 (PfBDP7) homologues and there is strong evidence for a role of this protein complex in maintaining mutually exclusive expression of a single surface antigen by repressing other surface antigen genes (18). Although we did not observe TgBDP1 bound at all *srs* gene loci that were upregulated during *tgbdp1* knockdown, it remains an intriguing possibility that the TgBDP1/2/5 complex contributes to a form of antigenic variation or virulence factor regulation in *Toxoplasma* by mediating stage-specific repression of these surface proteins.

TgBDP1 is recruited to over a thousand specific sites in the parasite genome. A large proportion of bindings sites correlate with both predicted TSSs (31) and histone marks linked with transcriptional initiation, but it is unclear how the TgBDP1 complex is recruited to these target binding sites. Interactome analysis identified a total of three bromodomain-containing proteins and a putative DNA binding protein (AP2VIIa-7) in the core complex indicating that this recruitment is mediated by recognition of a DNA sequence motif and/or histone modifications. Our analysis of the sequences around TgBDP1 TSS binding sites identified two significantly enriched sequence features, one of which, the GCATGC motif, has been reported previously in *Toxoplasma* (31,33,34) as an enriched motif at TSSs in tachyzoites, so is unlikely to be a specific motif to recruit the TgBDP1 complex and rather a general feature of *Toxoplasma* gene promoters. It is likely that interactions between the three bromodomains in the complex, and histone acetyl marks are more important mediators of complex recruitment to the chromatin. Determining the histone marks that are preferentially bound by the TgBDP1 bromodomain and the two interacting bromodomain-containing proteins TgBDP2 and TgBDP5 will be an important next step in understanding the function of this complex. We observed a strong correlation between TgBDP1 target sites and the histone H3K9Ac modification, suggesting that this may be one of the histone marks that is “read” by one or more of the bromodomains in the TgBDP1/2/5 complex. However, TgBDP1 was not located at every H3K9Ac enriched site indicating that another histone mark dictates TgBDP1/2/5 complex recruitment to the chromatin. The *P. falciparum* homologue, PfBDP1 did not display strong affinity for acetylated histone H3 peptide, even when multiple lysine residues were acetylated, suggesting that acetylated H3 is not important for complex recruitment. However, PfBDP1 has high binding affinity for acetylated histones H4, H2b.Z and H2a.z (24,44), particularly when multiple acetyl marks are present, suggesting that the PfBDP1/BDP2 complex is recruited to highly acetylated chromatin, rather than a specific acetylated histone residue. Additional studies will be needed to determine if this is also the case in *Toxoplasma*. Furthermore, it is probable that the ankyrin repeats of TgBDP1 mediate another important TgBDP1-protein interaction. The ankyrin repeats of the human methyltransferases G9a and G9a-like are reader modules of mono- and di-methylated H3K9 (45), so the ankyrin repeats of TgBDP1 may function in a similar manner to target the complex to modifications on chromatin. We also cannot discount the possibility that the bromodomain proteins may recognize a non-histone acetylated protein, of which many have been reported in *Toxoplasma* in previous studies (46– 48). The regulatory implications of acetylation marks on non-histone proteins in the nucleus have not been investigated, but it is likely that this process contributes to another level of transcriptional regulation.

Several aspects of TgBDP1 biological function parallel that of the *P. falciparum* homologue PfBDP1, including interacting closely with additional bromodomain-containing proteins, binding at TSSs, and regulating gene expression (16,18). However, we identified key differences that suggest distinct roles between species. Knockdown of PfBDP1 negatively impacted parasite invasion but with no effect on replication or overall fitness. Whereas TgBDP1 knockdown resulted in major defects to replication and ultimately parasite death. We found that TgBDP1 associates with a larger cohort of epigenetic regulators compared to the PfBDP1 associated proteins and regulates a larger proportion, and more functionally diverse sets of genes, suggesting TgBDP1 has additional functions beyond those reported for the *P. falciparum* homologue.

We show that while TgBDP1 has a highly conserved bromodomain, it is otherwise divergent from human, yeast, and plant bromodomain proteins. Only apicomplexans and a small number of other alveolates possess a homologous protein. As this essential protein is highly conserved amongst the human pathogens within the phylum Apicomplexa, it serves as a promising candidate for drug development that warrants more in-depth study.

## Methods

### Cell culture

*Toxoplasma gondii* tachyzoites of RHΔHXΔKu80 and TATiΔKu80 backgrounds (49,50)were maintained in human foreskin fibroblast (HFF) cells and Dulbecco’s Modified Eagle Medium supplemented with 1% fetal bovine serum. Cells were cultured in a humidified incubator at 37°C with 5% CO2. ^tet-myc^TgBDP1 and TgBDP1-HA cell lines were maintained in media containing 1uM pyrimethamine.

### TgBDP1 sequence analyses

The predicted TGME49_283580 (TgBDP1) genomic DNA, mRNA and protein sequences were obtained from ToxoDB (https://toxodb.org) (25). BLASTp analyses using the TgBDP1 predicted protein sequence were conducted in both the NCBI and VEuPathDB databases to identify homologues. Protein sequences from BDP1 homologues were aligned and evolutionary analysis performed using the maximum likelihood method in MEGA11 (51). Clustal Omega (EMBL-EBI) was used to align bromodomain sequences. The structure of TgBDP1’s bromodomain was predicted using I-TASSER and overlayed with the experimentally determined structures of the human B2AZB bromodomain (PDB 5DYU) and the *Plasmodium falciparum* PfBDP1 bromodomain (PDB 7M97) using Chimera (https://www.rbvi.ucsf.edu/chimerax) (52). The ToxoDB genome browser, JBrowse, was used to visualize predicted introns and nanopore mRNA sequencing reads from the Lee et al. dataset (26). To confirm TgBDP1 mRNA transcript sequences, RNA was harvested from RHΔHXΔKu80 parasites and cDNA synthesized using the Omniscript RT kit (Qiagen 205113) with *tgbdp1* specific primers 5’TTCAAAGATATGTCCACCCTCG and 5’CCTTACATCAGCAGACCTGC. The resulting cDNA was used to amplify tgbdp1 transcripts with primers 5’AGTGAATTCGAGCTCGGTACCATGTCGACTGGCGCGAGTG and 5’TGCATGCCTGCAGGTCGACTCTAGATTAAGCTCCACGTGATTCTCCG which were then cloned into a pUC19 vector. Plasmid DNA was isolated from six different bacterial clones and sequenced.

### Generation of TgBDP1 knockdown (^tet-myc^TgBDP1)

Inducible knockdown of the *tgbdp1* gene was accomplished by replacing the endogenous *tgbdp1* promoter with a tetracycline regulatable *tgsag4* promoter and adding a 3xmyc tag to the 5’end of the *tgbdp1* gene. A 2,100bp region of genomic DNA directly downstream of the *tgbdp1* start codon was amplified with primers 5’catctccgaggaggacctgagatctTCGACTGGCGCGAGTGTG and 5’TACGATGCGGCCGCcgatacatctgggcttgcc from TATiΔKu80 genomic DNA. The DHFR-tet07Sag4-3xMyc-CEP250 plasmid (kindly provided by MJ Gubbels) was digested with BglII and NotI, and the HiFi DNA Assembly kit (NEB E5520S) was used to insert the *tgbdp1* fragment. The final DHFR-tet07Sag4-3xMyc-tgbdp1 plasmid was verified by sequencing. One hundred micrograms of plasmid was linearized with NotI and transfected into TATiΔKu80 (TATi) parasites (kindly provided by MJ Gubbels) by electroporation in cytomix (53). Parasites were cultured with 1uM pyrimethamine for selection and cloned by limiting dilution. PCR of genomic DNA with primers P1 5’GCTAATCTCCGAGGAAGACTTG and P2 5’TGGCCTGCTCTCGTTTCAC was used to confirm correct integration. PCR with primers P2 5’CGATTGCCTCTCCCTCAAGTCC and P3 5’TCTCGACCTCTTCGCGTACG confirmed disruption of the endogenous promoter.

### Generation of endogenously tagged TgBDP1 (TgBDP1^3xHA^)

A 3xHA epitope tag was introduced at the 3’end of the endogenous *tgbdp1* gene. A 2,131bp region of genomic DNA upstream of the *tgbdp1* stop codon was amplified with primers 5’ tacttccaatccaatttaattaaTGAGCAAGTGAGGCAAGC and 5’cctccacttccaattttaattaaAGCTCCACGTGATTCTCC and inserted using the HiFi DNA assembly kit into pLIC-3xHA-DHFR cut with PacI. The final pLIC-TgBDP1-3xHA-DHFR plasmid was verified by sequencing. One hundred micrograms of plasmid was linearized with AflII and transfected into RHΔHXΔKu80 (ΔKu80) parasites by electroporation in cytomix. Parasites were cultured with 1uM pyrimethamine for selection and cloned by limiting dilution.

### Immunofluorescence Assays

Parasites were inoculated into 24-well plates of confluent HFFs containing coverslips and cultured approximately 24hrs (+/-ATc), then fixed with 4% paraformaldehyde and permeabilized with Triton X-100 in 3% BSA. Samples were blocked in 3% BSA and primary and secondary antibodies diluted in 3% BSA. Primary antibodies anti-myc (Invitrogen 132500) and anti-HA (Roche 27573500) were diluted 1:2000 and secondary antibodies anti-mouse Alexa Fluor 594 (Thermo Fisher Scientific A11005) and anti-rat Alexa Fluor 594 (Thermo Fisher Scientific A11007) were diluted 1:5000. After antibody incubations, samples were incubated with DAPI (Invitrogen D1306) in 3% BSA and coverslips mounted to slides with Vectashield mounting medium (Vector Laboratories H1000). Slides were visualized and imaged with a Nikon A1R laser scanning Confocal Fluorescence Microscope and NIS-Elements software.

### Western blotting

Protein was isolated from parasites by resuspending harvested parasites in RIPA buffer supplemented with protease inhibitor cocktail (Research Products International Corp P506001). Samples were then sonicated using a QSonica Q800R3 at 50% amplitude for 2 minutes. Insoluble material was pelleted and removed. Protein concentrations of lysates were determined using a BCA kit (Thermo Fisher Scientific 23227) and 50ug of protein was used for Western blotting. Protein samples were separated by SDS-PAGE in 4-15% Bis-Tris gels with MOPS buffer and transferred to nitrocellulose membrane. Membranes were blocked in 5% non-fat milk and incubated in primary and secondary antibodies diluted in 5% non-fat milk. The following antibodies were used: anti-myc-HRP (Santa Cruz sc-40) diluted 1:100, anti-HA (Roche 27573500) diluted 1:2000, anti-rat-HRP (GE NA935) diluted 1:2000, anti-p30 (SAG1) (Invitrogen MA183499) diluted 1:2000 and anti-mouse-HRP (GE NA931) diluted 1:2000. Pierce ECL detection reagent (Thermo Fisher Scientific 32109) and a BioRadV3 Chemidoc Imager were used to visualize blots.

### Quantitative RT-PCR

Total RNA was harvested from ^tet-myc^TgBDP1 parasites 36hrs post-inoculation and that had been cultured +/-ATc 1uM ATc for 12, 24 or 36 hours. Parasites were pelleted and resuspended in 1ml TRI Reagent (MilliporeSigma T9424). RNA was isolated by phenol:chloroform extraction and isopropanol precipitation followed by genomic DNA removal using the Turbo DNA-free kit (Invitrogen AM1907). Three micrograms of RNA were used to synthesize cDNA with the Omniscript RT kit using oligo dT primers (Qiagen 205113). Resulting cDNA was diluted 1:2 and used for real-time PCR with Power SYBR Green (Thermo Fisher Scientific 4367659) and Applied Biosystems 7500 real time PCR system. *tgbdp1* was amplified with primers 5’CACATCCTCAGCAATTCCTTAAG and 5’GCGAGGACACTGTAGATCTTG, and *tgtuba1* (used for normalization) was amplified with primers 5’GATGCCCTCTGACAAGACC and 5’CATCCTCTTTCCCGCTGATC. The delta delta Ct method was used to quantify changes in gene expression compared to -ATc samples and 2^-ddct was used to calculate fold change. Data from three independent replicates was statistically analyzed using one-way ANOVA and Dunnett’s multiple comparisons test in GraphPad Prism Version 9.3.1 for MacOS, GraphPad Software, San Diego, California USA, www.graphpad.com.

### Toxoplasma growth assays

Plaque assays were done to assess the effect of TgBDP1 knockdown on *Toxoplasma* growth as previously described (53). Briefly, 200 parasites of the ^tet-myc^TgBDP1 and parental (TATi) parasite lines were inoculated into 12-well plates of confluent HFFs in media +/-ATc and cultured for six days. Cells were then fixed in methanol, stained with crystal violet, and imaged with an Invitrogen EVOS M7000 microscope. The area of the plaques per well (area of host cell lysis) was quantified from the images using ImageJ software and percentage of host cell lysis compared to -ATc calculated. An unpaired t-test from three independent experiments was performed using GraphPad Prism.

*Toxoplasma* red/green invasion assays were performed as previously described (54). ^tet-myc^TgBDP1 and TATi parasites were cultured +/-ATc for 36hrs, at which point intracellular parasites were harvested and counted. Parasites and 12-well plates of HFFs containing coverslips were chilled on ice and 1×10^6 parasites inoculated per well, remaining on ice for 15min. The inoculated plate was then incubated in a 37°C water bath for 1min before moving to the 37°C incubator. Plates were incubated for 2hr, then washed to remove extracellular parasites. Cells were fixed with 3% paraformaldehyde, blocked with 3% BSA and incubated with 1:1000 dilution of mouse anti-P30 (SAG1) primary antibody (Invitrogen MA183499). Cells were then permeabilized and incubated with 1:1000 dilution of rabbit anti-Toxoplasma primary antibody (Invitrogen PA17252) followed by a final incubation with secondary antibodies goat anti-mouse Alexa Fluor 488 (1:5000) (Thermo Fisher Scientific A11001) and goat anti-rabbit Alexa Flour 594 (1:5000) (Thermo Fisher Scientific A11012). A Zeiss Axioplan 2 fluorescent microscope was used to visualize over 1000 parasites per treatment group. Red only parasites were designated intracellular while dual color parasites (red and green) were considered extracellular. The percentage of intracellular parasites was calculated and an unpaired t-test between -ATc and +ATc groups for three independent experiments performed.

Doubling, or replication, assays were used to determine *Toxoplasma* replication rate. ^tet-myc^TgBDP1 and TATi parasites were cultured +/-ATc for 24hrs, at which point intracellular parasites were harvested and inoculated into a 12-well plate of confluent HFFs in media +/-ATc. Two hours post-inoculation media and extracellular parasites were removed and fresh media +/-ATc added. Wells were fixed with Hema3 fixative 12, 24 and 36hrs post-inoculation and then stained with Hema3 Staining Solutions I and II. The number of parasites per vacuole was counted for 100 vacuoles. Three independent experiments were conducted.

### Co-immunoprecipitation

CoIPs and mass spectrometry were used to identify TgBDP1-interacting proteins in TgBDP1-HA parasites, with ΔKu80 parasites used as a negative control. For each sample, parasites were cultured for 36hrs and 8 T-150s of intracellular parasites harvested. Nuclei were harvested by resuspending cells in 1ml lysis buffer A [10mM KCl, 10mM HEPES pH 7.4, 0.1% NP-40, 10% glycerol, cOmplete protease inhibitor cocktail (Roche 04693159001)], incubated on ice 5min then pelleted at 10,000xg for 10min at 4°C. The nuclei pellet was resuspended in 500ul lysis buffer B (400mM KCl, 10mM HEPES pH 7.4, 0.1% NP-40, 10% glycerol, cOmplete protease inhibitor cocktail), vortexed for 30min at 4°C and centrifuged at 10,000xg for 10min at 4°C. For each 500ul nuclear supernatant, 50ul of pre-washed anti-HA magnetic beads (Thermo Fisher Scientific 88837) was added and samples rocked at 4°C overnight. Protein-bound anti-HA magnetic beads were washed five times in coIP buffer (0.025M Tris, 0.15M NaCl, 0.001M EDTA, 1% NP-40, 5% glycerol, cOmplete protease inhibitor cocktail) then resuspended in 45ul 2x LDS buffer and 8% beta-mercaptoethanol and boiled 10min. Samples were run on a 4-12% Bis-Tis gel in MOPS buffer and the gel stained with Coomassie G-250 for 1.5hrs.

Protein samples were recovered by isolating gel bands that were then processed via in-gel digestion and analyzed by LC-MS and LC-MS-MS as described previously (55). Briefly, a 1 ul aliquot of the digestion mixtures was injected into a Dionex Ultimate 3000 RSLCnano UHPLC system with an autosampler (Dionex Corporation, Sunnyvale, CA, USA), where it was then separated in a 100 µm x 15 cm capillary packed with Dr. Maisch ReproSil-Pur C18-AQ, r13.aq (120 Å; 3 µm), at a flow rate of ∼450 nl/min. The eluant was connected directly to a nanoelectrospray ionization source of an LTQ Orbitrap XL mass spectrometer (ThermoFisher Scientific). LC-MS data were acquired in a data-dependent acquisition mode, cycling between a MS scan (m/z 315-2000) acquired in the Orbitrap, followed by collision-induced dissociation analysis on the three most intensely multiply charged precursors acquired in the linear ion trap. The LC-MS/MS data was processed by PAVA bioinformatic program to generate the centroided peak lists of the CID spectra and searched against a database that consisted of the Swiss-Prot protein database (version 2021.06.18; 53/565,254 entries searched for Toxoplasma Gondii), using the Batch-Tag program module of the Protein Prospector bioinformatic package from the University of California, San Francisco (version 6.3.1). A precursor mass tolerance of 20 ppm and a fragment mass tolerance of 0.6 Da were used for protein database search (trypsin as enzyme; one missed cleavage; carbamidomethyl [C] as constant modification; acetyl [protein N-term], acetyl + oxidation [protein N-term M], Gln - >pyro-Glu [N-term Q], Met-loss [protein N-term M], Met-loss +acetyl [protein N-term M], oxidation [M] as variable modifications). Protein matches were reported with a Protein Prospector protein score ≥22, a protein discriminant score ≥0.0, and a peptide expectation value ≤0.01 (56). Data are available via ProteomeXchange with identifier PXD038848. Potential TgBDP1-associated proteins were identified as those with peptide counts at least 2-fold higher in TgBDP1-HA samples compared to ΔKu80, present in at least 2 out of 3 independent experiments and with a predicted or verified nuclear localization. SAINT analysis was performed using REPRINT (https://reprint-apms.org/) with default settings (28).

### Cleavage Under Targets & Tagmentation (CUT&Tag)

CUT&Tag was used to identify the genomic localization of TgBDP1^3xHA^. Based on protocols and findings from the Henikoff lab (29), we modified the technique for use with *Toxoplasma* tachyzoites. For each sample, intracellular parasites cultured for 36hrs were harvested from 1 T-75, syringe lysed, filtered through a 3um filter and counted. Ten million (1×10^7) parasites were centrifuged 2000xg for 10min and the parasite pellet used directly with the CUT&Tag-IT Assay Kit (Active Motif 53160). Our experimental sample used TgBDP1^3xHA^ parasites and 1ul of a 1:50 dilution of the rabbit anti-HA primary antibody (Cell Signaling 3724T). The negative control sample used ΔKu80 parasites (the parental line) with the same primary antibody conditions. Samples were incubated with primary antibody overnight at 4°C. The remainder of the kit protocol was followed exactly, and unique indexed primers were used for each sample. Three biological replicates were done for both TgBDP1^3xHA^ and ΔKu80 parasites. Due to small cell number and low amount of input DNA needed for this technique, negative controls often result in very little to no DNA and are therefore not able to be sequenced. This was the case for one or our three negative control replicates. A single positive control sample was processed in parallel to confirm that our technique was successful. We used an anti-H3K9ac antibody (Active Motif 39917) with TgBDP1^3xHA^ parasites to identify the highly abundant H3K9ac mark throughout the *Toxoplasma* genome, which has previously been done using ChIP-chip (30).

Indexed libraries for each sample were analyzed by TapeStation, pooled and run on NextSeq 500/550 High Output (75 cycles) flow cell to generate paired end reads. Demultiplexing of the reads was performed with bcl2fastq version 2.20.0 and processed with cutadapt v3.4 (57) to filter sequencing adapters from the 3’ end of reads, and any reads with fewer than 30 base pairs were removed. Remaining reads were mapped to version 52 of the Toxoplasma gondii ME49 reference downloaded from ToxoDB (25). The mapping was performed with HISAT2 v 2.2.1 (58) using the following parameters “--no-discordant--no-spliced-alignment --phred33 --no-unal --nomixed”. Unmapped reads were removed with samtools 1.9 (59). For each sample, peaks were then called using the callpeak command within MACS3 3.0.0a6 (60), with the parameters “-g 6.e7 -B -q 0.01”. A custom python script was used to associate predicted peaks with genes defined in version 52 of the ME49 reference and Transcription Start Sites identified in *Toxoplasma gondii* (31). A heatmap of mapped reads in relation to these TSS’s was generated using the computeMatrix and plotHeatmap commands from deepTools version 3.5.0 (61). DNA motif analysis was performed using MEME-ChIP (62) with default settings and sequences 250bp upstream and downstream of TSS with TgBDP1 associated peaks.

### RNA-sequencing (RNA-seq)

^tet-myc^TgBDP1 parasites were cultured 36hrs and treated -ATc or +ATc for 12, 24 or 36hrs. Intracellular parasites were harvested from 2 T-150s for each sample, syringe lysed, filtered through a 3um filter and pelleted. The Qiagen RNease Plus Mini Kit was used to isolate RNA per the manufacturer’s instructions. Library preparation was completed with the KAPA mRNA HyperPrep Kit (Illumina® Platforms). Sequencing was completed at the Hubbard Center for Genome Studies on an Illumina NovaSeq 6000 platform to produce 250bp paired-end reads. Raw sequencing data was demultiplexed using bcl2fastq v1.8.4 (Illumina). Read quality was examined with FASTQC v0.11.9. Adapters and low-quality sequences were trimmed from the reads using Trimmomatic V0.32 (63) with default setting. The *Toxoplasma* reference genome and annotations (ME49) were downloaded from ToxoDB (release 56), and datasets were mapped to the reference genome using HISAT2 (58) with default setting. The counts of reads mapping to each gene feature in the GFF annotations was completed using featureCounts (64). The outputs from featureCounts were analyzed within RStudio (Build 443) following the DESeq2 v1.32.0 vignette (65).

## Acknowledgements

The authors are grateful to Dr. Marc Jan Gubbels (Boston College) for sharing plasmids; Dr. Kelley Thomas, Steven Smith and Joe Sevigny at the UNH Hubbard Center for Genomics for assistance with RNA sequencing and analysis; and Dr. Doug Rusch and Chris Hemmerich at IU Center for Genomics and Bioinformatics for assistance with CUT&Tag sequencing and analysis. Molecular graphics and analyses performed with UCSF Chimera, developed by the Resource for Biocomputing, Visualization, and Informatics at the University of California, San Francisco, with support from NIH P41-GM103311. VJ is supported as a Project Lead by CIBBR through a grant from NIGMS (P20GM113131) at NIH.

---

**Supplemental Figure 1.**
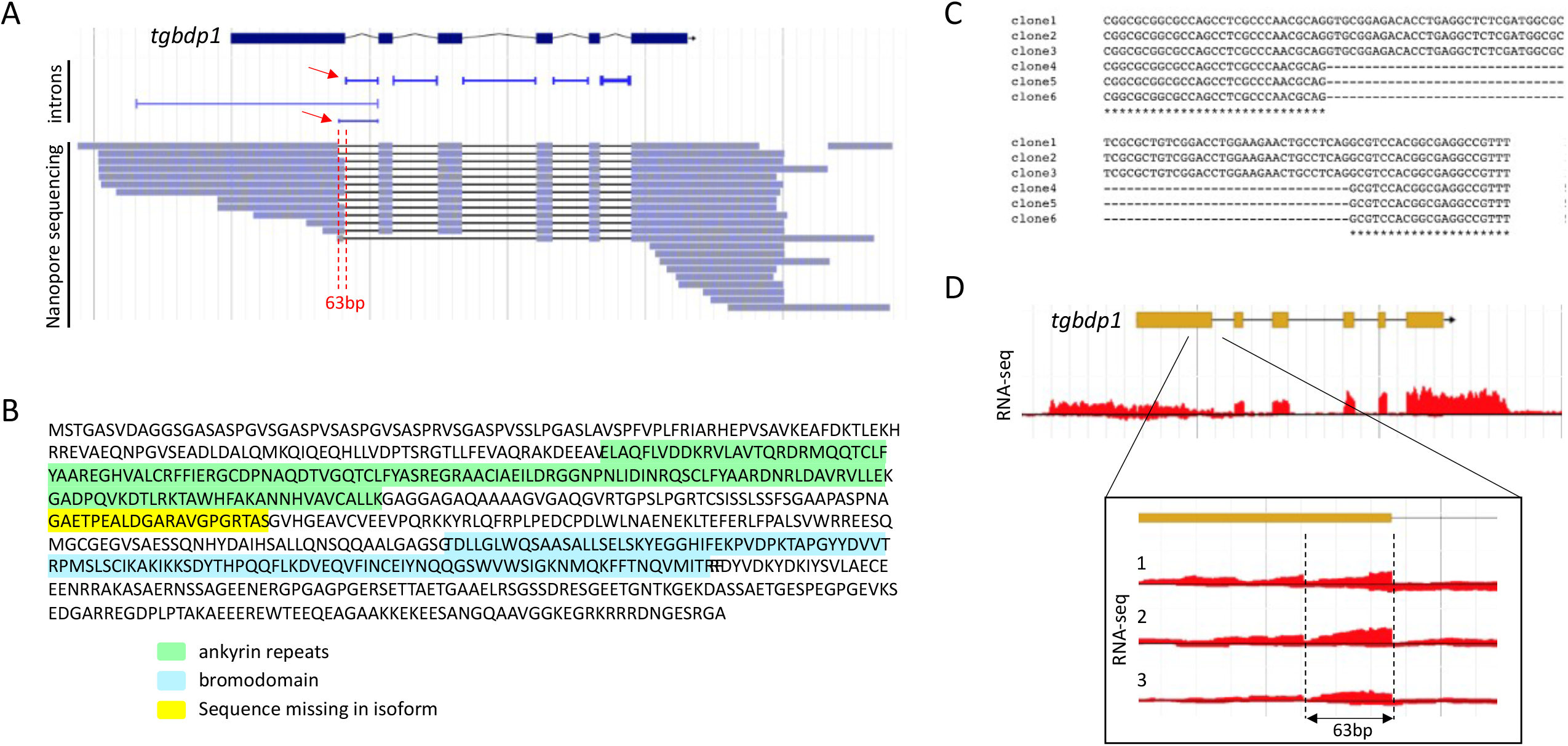
TgBDP1 has an mRNA isoform, TgBDP1a. A) Screen capture from ToxoDB genome browser showing the predicted tgbdp1 gene with exons (blue boxes) and intron (lines) on top. Underneath is predicted introns with red arrows identifying the two distinct isoforms. Nanopore sequencing read alignments of *Toxoplasma* mRNAs are shown with the 63 nucleotide isoform region flanked by red dashed lines. B) Predicted TgBDP1 protein sequence with ankyrin repeats highlighted in green, bromodomain in blue and the sequence missing from TgBDP1a in yellow. C) Multiple sequence alignment of cDNA encompassing the 63 isoform nucleotides from six different *tgbdp1* clones. Clones 1-3 have the full predicted sequence (*tgbdp1*) while clones 3-6 are missing the 63 nucleotides (*tgbdp1a*). D) Screen capture from ToxoDB genome browser showing the *tgbdp1* gene model with exons (yellow boxes) and introns (lines), with RNA-sequencing peaks of one replicate from our parasite line ^tet-myc^TgBDP1. Inset depicts the end of the first exon and beginning of the first intron, and RNA-sequencing peaks from all three replicates with the 63 nucleotide region flanked by dashed lines.

**Supplemental Figure 2.**
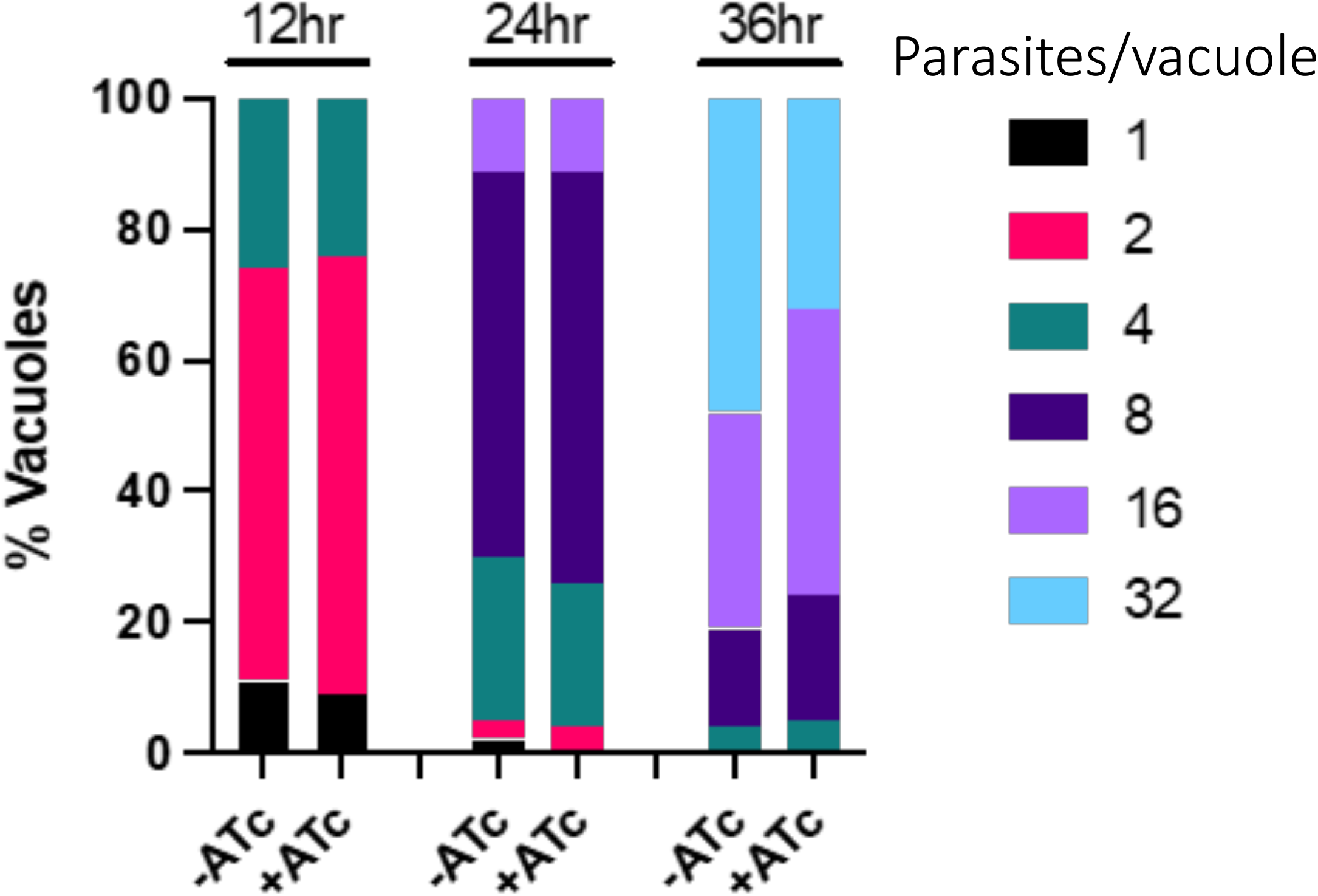
Parental parasite line (TATi) replicates normally in the presence of ATc. A *Toxoplasma* doubling assay was performed with parasites -ATc and +ATc. The number of parasites per vacuole was counted at 12, 24 and 36hrs after inoculation.

**Supplemental Figure 3.**
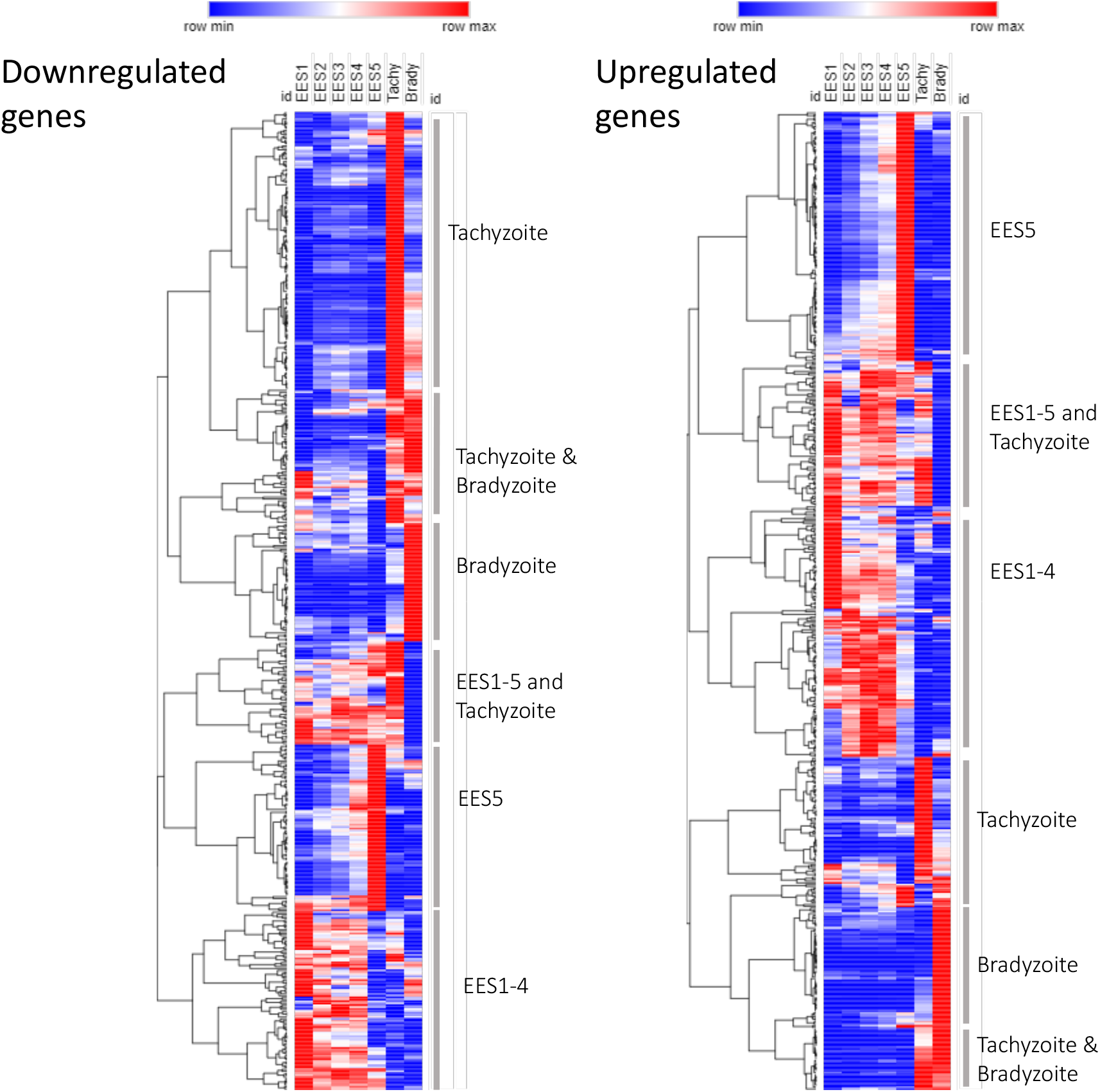
Life cycle expression of genes impacted during *tgbdp1* knockdown in tachyzoites. Heat maps of relative gene expression levels of the genes that are significantly downregulated (left) and upregulated (right) during *tgbdp1* knockdown. Gene expression data for parasite stages from Ramakrishnan *et al*. (41) was clustered according to the life cycle stage(s) in which expression peaks. Labels indicate the stage of peak gene expression for each gene cluster.

## References

1. Dixon SE, Stilger KL, Elias EV, Naguleswaran A, Sullivan WJ. A decade of epigenetic research in Toxoplasma gondii. Mol Biochem Parasitol. 2010 Sep;173(1):1–9.

2. Vanagas L, Jeffers V, Bogado SS, Dalmasso MC, Sullivan WJ, Angel SO. Toxoplasma histone acetylation remodelers as novel drug targets. Expert Rev Anti Infect Ther. 2012 Oct;10(10):1189–201.

3. Nardelli SC, Che FY, Monerri NCS de, Xiao H, Nieves E, Madrid-Aliste C, et al. The Histone Code of Toxoplasma gondii Comprises Conserved and Unique Posttranslational Modifications. mBio [Internet]. 2013 Dec 31 [cited 2020 Aug 20];4(6). Available from: https://mbio.asm.org/content/4/6/e00922-13

4. Bhatti MM, Livingston M, Mullapudi N, Sullivan WJ. Pair of unusual GCN5 histone acetyltransferases and ADA2 homologues in the protozoan parasite Toxoplasma gondii. Eukaryot Cell. 2006 Jan;5(1):62–76.

5. Harris MT, Jeffers V, Martynowicz J, True JD, Mosley AL, Sullivan WJ. A novel GCN5b lysine acetyltransferase complex associates with distinct transcription factors in the protozoan parasite Toxoplasma gondii. Mol Biochem Parasitol. 2019 Sep 1;232:111203.

6. Naguleswaran A, Elias EV, McClintick J, Edenberg HJ, Sullivan WJ. Toxoplasma gondii lysine acetyltransferase GCN5-A functions in the cellular response to alkaline stress and expression of cyst genes. PLoS Pathog. 2010 Dec 16;6(12):e1001232.

7. Wang J, Dixon SE, Ting LM, Liu TK, Jeffers V, Croken MM, et al. Lysine acetyltransferase GCN5b interacts with AP2 factors and is required for Toxoplasma gondii proliferation. PLoS Pathog. 2014 Jan;10(1):e1003830.

8. Sindikubwabo F, Ding S, Hussain T, Ortet P, Barakat M, Baumgarten S, et al. Modifications at K31 on the lateral surface of histone H4 contribute to genome structure and expression in apicomplexan parasites. Zilberman D, editor. eLife. 2017 Nov 4;6:e29391.

9. Bougdour A, Maubon D, Baldacci P, Ortet P, Bastien O, Bouillon A, et al. Drug inhibition of HDAC3 and epigenetic control of differentiation in Apicomplexa parasites. J Exp Med. 2009 Apr 6;206(4):953–66.

10. Saksouk N, Bhatti MM, Kieffer S, Smith AT, Musset K, Garin J, et al. Histone-Modifying Complexes Regulate Gene Expression Pertinent to the Differentiation of the Protozoan Parasite Toxoplasma gondii. Mol Cell Biol. 2005 Dec;25(23):10301–14.

11. Hanquier J, Gimeno T, Jeffers V, Sullivan WJ. Evaluating the GCN5b bromodomain as a novel therapeutic target against the parasite Toxoplasma gondii. Exp Parasitol. 2020 Apr 1;211:107868.

12. Jeffers V, Gao H, Checkley LA, Liu Y, Ferdig MT, Sullivan WJ. Garcinol Inhibits GCN5-Mediated Lysine Acetyltransferase Activity and Prevents Replication of the Parasite Toxoplasma gondii. Antimicrob Agents Chemother. 2016 Apr;60(4):2164–70.

13. Darkin-Rattray SJ, Gurnett AM, Myers RW, Dulski PM, Crumley TM, Allocco JJ, et al. Apicidin: A novel antiprotozoal agent that inhibits parasitelJhistonelJdeacetylase. Proc Natl Acad Sci. 1996 Nov 12;93(23):13143–7.

14. Dhalluin C, Carlson JE, Zeng L, He C, Aggarwal AK, Zhou MM. Structure and ligand of a histone acetyltransferase bromodomain. Nature. 1999 Jun;399(6735):491–6.

15. Kulikowski E, Rakai BD, Wong NCW. Inhibitors of bromodomain and extra-terminal proteins for treating multiple human diseases. Med Res Rev. 2021;41(1):223–45.

16. Josling GA, Petter M, Oehring SC, Gupta AP, Dietz O, Wilson DW, et al. A Plasmodium Falciparum Bromodomain Protein Regulates Invasion Gene Expression. Cell Host Microbe. 2015 Jun 10;17(6):741–51.

17. Santos JM, Josling G, Ross P, Joshi P, Orchard L, Campbell T, et al. Red Blood Cell Invasion by the Malaria Parasite Is Coordinated by the PfAP2-I Transcription Factor. Cell Host Microbe. 2017 Jun 14;21(6):731-741.e10.

18. Quinn JE, Jeninga MD, Limm K, Pareek K, Meißgeier T, Bachmann A, et al. The Putative Bromodomain Protein PfBDP7 of the Human Malaria Parasite Plasmodium Falciparum Cooperates With PfBDP1 in the Silencing of Variant Surface Antigen Expression. Front Cell Dev Biol [Internet]. 2022 [cited 2022 Nov 30];10. Available from: https://www.frontiersin.org/articles/10.3389/fcell.2022.816558

19. Jeffers V, Kamau ET, Srinivasan AR, Harper J, Sankaran P, Post SE, et al. TgPRELID, a Mitochondrial Protein Linked to Multidrug Resistance in the Parasite Toxoplasma gondii. mSphere. 2017 Feb;2(1).

20. Jeffers V, Yang C, Huang S, Sullivan WJ. Bromodomains in Protozoan Parasites: Evolution, Function, and Opportunities for Drug Development. Microbiol Mol Biol Rev MMBR. 2017 Mar;81(1).

21. Fleck K, Nitz M, Jeffers V. “Reading” a new chapter in protozoan parasite transcriptional regulation. PLOS Pathog. 2021 Dec 2;17(12):e1010056.

22. Owen DJ, Ornaghi P, Yang JC, Lowe N, Evans PR, Ballario P, et al. The structural basis for the recognition of acetylated histone H4 by the bromodomain of histone acetyltransferase Gcn5p. EMBO J. 2000 Nov 15;19(22):6141–9.

23. Yang J, Yan R, Roy A, Xu D, Poisson J, Zhang Y. The I-TASSER Suite: protein structure and function prediction. Nat Methods. 2015 Jan;12(1):7–8.

24. Singh AK, Phillips M, Alkrimi S, Tonelli M, Boyson SP, Malone KL, et al. Structural insights into acetylated histone ligand recognition by the BDP1 bromodomain of Plasmodium falciparum. Int J Biol Macromol. 2022 Dec 31;223:316–26.

25. Amos B, Aurrecoechea C, Barba M, Barreto A, Basenko EY, Bażant W, et al. VEuPathDB: the eukaryotic pathogen, vector and host bioinformatics resource center. Nucleic Acids Res. 2022 Jan 7;50(D1):D898–911.

26. Lee VV, Judd LM, Jex AR, Holt KE, Tonkin CJ, Ralph SA. Direct Nanopore Sequencing of mRNA Reveals Landscape of Transcript Isoforms in Apicomplexan Parasites. mSystems. 2021 Mar 9;6(2):e01081–20.

27. Sidik SM, Huet D, Ganesan SM, Huynh MH, Wang T, Nasamu AS, et al. A Genome-wide CRISPR Screen in Toxoplasma Identifies Essential Apicomplexan Genes. Cell. 2016 Sep 8;166(6):1423-1435.e12.

28. Mellacheruvu D, Wright Z, Couzens AL, Lambert JP, St-Denis NA, Li T, et al. The CRAPome: a contaminant repository for affinity purification-mass spectrometry data. Nat Methods. 2013 Aug;10(8):730–6.

29. Kaya-Okur HS, Wu SJ, Codomo CA, Pledger ES, Bryson TD, Henikoff JG, et al. CUT&Tag for efficient epigenomic profiling of small samples and single cells. Nat Commun. 2019 Dec;10(1):1930.

30. Gissot M, Kelly KA, Ajioka JW, Greally JM, Kim K. Epigenomic Modifications Predict Active Promoters and Gene Structure in Toxoplasma gondii. PLOS Pathog. 2007 Jun 8;3(6):e77.

31. Markus BM, Waldman BS, Lorenzi HA, Lourido S. High-Resolution Mapping of Transcription Initiation in the Asexual Stages of Toxoplasma gondii. Front Cell Infect Microbiol. 2020;10:617998.

32. Waldman BS, Schwarz D, Wadsworth MH, Saeij JP, Shalek AK, Lourido S. Identification of a Master Regulator of Differentiation in Toxoplasma. Cell. 2020 Jan 23;180(2):359-372.e16.

33. Behnke MS, Wootton JC, Lehmann MM, Radke JB, Lucas O, Nawas J, et al. Coordinated Progression through Two Subtranscriptomes Underlies the Tachyzoite Cycle of Toxoplasma gondii. PLOS ONE. 2010 Aug 26;5(8):e12354.

34. Van Poppel NFJ, Welagen J, Vermeulen AN, Schaap D. The complete set of Toxoplasma gondii ribosomal protein genes contains two conserved promoter elements. Parasitology. 2006 Jul;133(Pt 1):19–31.

35. Yamagishi J, Wakaguri H, Ueno A, Goo YK, Tolba M, Igarashi M, et al. High-Resolution Characterization of Toxoplasma gondii Transcriptome with a Massive Parallel Sequencing Method. DNA Res. 2010 Aug 1;17(4):233–43.

36. Shea M, Jäkle U, Liu Q, Berry C, Joiner KA, Soldati-Favre D. A family of aspartic proteases and a novel, dynamic and cell-cycle-dependent protease localization in the secretory pathway of Toxoplasma gondii. Traffic Cph Den. 2007 Aug;8(8):1018–34.

37. Farhat DC, Swale C, Dard C, Cannella D, Ortet P, Barakat M, et al. A MORC-driven transcriptional switch controls Toxoplasma developmental trajectories and sexual commitment. Nat Microbiol. 2020 Apr;5(4):570–83.

38. Mineo TWP, Chern JH, Thind AC, Mota CM, Nadipuram SM, Torres JA, et al. Efficient Gene Knockout and Knockdown Systems in Neospora caninum Enable Rapid Discovery and Functional Assessment of Novel Proteins. mSphere. 2022 Jan 12;7(1):e00896–21.

39. Radke JB, Lucas O, De Silva EK, Ma Y, Sullivan WJ, Weiss LM, et al. ApiAP2 transcription factor restricts development of the Toxoplasma tissue cyst. Proc Natl Acad Sci U S A. 2013 Apr 23;110(17):6871–6.

40. Licon MH, Giuliano CJ, Chakladar S, Shallberg L, Waldman BS, Hunter CA, et al. A positive feedback loop controls Toxoplasma chronic differentiation [Internet]. bioRxiv; 2022 [cited 2022 Dec 1]. p. 2022.04.06.487076. Available from: https://www.biorxiv.org/content/10.1101/2022.04.06.487076v1

41. Ramakrishnan C, Maier S, Walker RA, Rehrauer H, Joekel DE, Winiger RR, et al. An experimental genetically attenuated live vaccine to prevent transmission of Toxoplasma gondii by cats. Sci Rep. 2019 Feb 6;9(1):1474.

42. Zhang Y, Cheng L, Qiu H, Sun T, Deng R, Gong H, et al. Hypothetical bromodomain-containing protein 5 is required for the growth of Toxoplasma gondii. Vet Parasitol. 2022 Sep 1;309:109767.

43. Theisen TC, Boothroyd JC. Transcriptional signatures of clonally derived Toxoplasma tachyzoites reveal novel insights into the expression of a family of surface proteins. PloS One. 2022;17(2):e0262374.

44. Hoeijmakers WAM, Miao J, Schmidt S, Toenhake CG, Shrestha S, Venhuizen J, et al. Epigenetic reader complexes of the human malaria parasite, Plasmodium falciparum. Nucleic Acids Res. 2019 Dec 16;47(22):11574–88.

45. Collins RE, Northrop JP, Horton JR, Lee DY, Zhang X, Stallcup MR, et al. The ankyrin repeats of G9a and GLP histone methyltransferases are mono- and dimethyllysine binding modules. Nat Struct Mol Biol. 2008 Mar;15(3):245–50.

46. Jeffers V, Sullivan WJ. Lysine acetylation is widespread on proteins of diverse function and localization in the protozoan parasite Toxoplasma gondii. Eukaryot Cell. 2012 Jun;11(6):735–42.

47. Xue B, Jeffers V, Sullivan WJ, Uversky VN. Protein intrinsic disorder in the acetylome of intracellular and extracellular Toxoplasma gondii. Mol Biosyst. 2013 Apr 5;9(4):645–57.

48. Kloehn J, Oppenheim RD, Siddiqui G, De Bock PJ, Kumar Dogga S, Coute Y, et al. Multi-omics analysis delineates the distinct functions of sub-cellular acetyl-CoA pools in Toxoplasma gondii. BMC Biol. 2020 Jun 16;18(1):67.

49. Sheiner L, Demerly JL, Poulsen N, Beatty WL, Lucas O, Behnke MS, et al. A systematic screen to discover and analyze apicoplast proteins identifies a conserved and essential protein import factor. PLoS Pathog. 2011 Dec;7(12):e1002392.

50. Huynh MH, Carruthers VB. Tagging of endogenous genes in a Toxoplasma gondii strain lacking Ku80. Eukaryot Cell. 2009 Apr;8(4):530–9.

51. Tamura K, Stecher G, Kumar S. MEGA11: Molecular Evolutionary Genetics Analysis Version 11. Mol Biol Evol. 2021 Jun 25;38(7):3022–7.

52. Pettersen EF, Goddard TD, Huang CC, Couch GS, Greenblatt DM, Meng EC, et al. UCSF Chimera--a visualization system for exploratory research and analysis. J Comput Chem. 2004 Oct;25(13):1605–12.

53. Jacot D, Frénal K, Marq JB, Sharma P, Soldati-Favre D. Assessment of phosphorylation in Toxoplasma glideosome assembly and function. Cell Microbiol. 2014 Oct;16(10):1518–32.

54. Suarez C, Lodoen MB, Lebrun M. Assessing Rhoptry Secretion in T. gondii. Methods Mol Biol Clifton NJ. 2020;2071:143–55.

55. Wu T, Nance J, Chu F, Fazzio TG. Characterization of R-Loop-Interacting Proteins in Embryonic Stem Cells Reveals Roles in rRNA Processing and Gene Expression. Mol Cell Proteomics MCP. 2021;20:100142.

56. Chalkley RJ, Baker PR, Huang L, Hansen KC, Allen NP, Rexach M, et al. Comprehensive analysis of a multidimensional liquid chromatography mass spectrometry dataset acquired on a quadrupole selecting, quadrupole collision cell, time-of-flight mass spectrometer: II. New developments in Protein Prospector allow for reliable and comprehensive automatic analysis of large datasets. Mol Cell Proteomics MCP. 2005 Aug;4(8):1194–204.

57. Martin M. Cutadapt removes adapter sequences from high-throughput sequencing reads. EMBnet.journal. 2011 May 2;17(1):10–2.

58. Kim D, Paggi JM, Park C, Bennett C, Salzberg SL. Graph-based genome alignment and genotyping with HISAT2 and HISAT-genotype. Nat Biotechnol. 2019 Aug;37(8):907–15.

59. Danecek P, Bonfield JK, Liddle J, Marshall J, Ohan V, Pollard MO, et al. Twelve years of SAMtools and BCFtools. GigaScience. 2021 Feb 16;10(2):giab008.

60. Zhang Y, Liu T, Meyer CA, Eeckhoute J, Johnson DS, Bernstein BE, et al. Model-based Analysis of ChIP-Seq (MACS). Genome Biol. 2008 Sep 17;9(9):R137.

61. Ramírez F, Ryan DP, Grüning B, Bhardwaj V, Kilpert F, Richter AS, et al. deepTools2: a next generation web server for deep-sequencing data analysis. Nucleic Acids Res. 2016 Jul 8;44(W1):W160–165.

62. Machanick P, Bailey TL. MEME-ChIP: motif analysis of large DNA datasets. Bioinforma Oxf Engl. 2011 Jun 15;27(12):1696–7.

63. Bolger AM, Lohse M, Usadel B. Trimmomatic: a flexible trimmer for Illumina sequence data. Bioinforma Oxf Engl. 2014 Aug 1;30(15):2114–20.

64. Liao Y, Smyth GK, Shi W. featureCounts: an efficient general purpose program for assigning sequence reads to genomic features. Bioinforma Oxf Engl. 2014 Apr 1;30(7):923–30.

65. Love MI, Huber W, Anders S. Moderated estimation of fold change and dispersion for RNA-seq data with DESeq2. Genome Biol. 2014;15(12):550.

